# Dopamine-dependent cerebellar dysfunction enhances beta oscillations and disrupts motor learning in a multiarea model

**DOI:** 10.1101/2023.07.18.549459

**Authors:** Benedetta Gambosi, Francesco Jamal Sheiban, Marco Biasizzo, Alberto Antonietti, Egidio D’Angelo, Alberto Mazzoni, Alessandra Pedrocchi

**Affiliations:** NearLab, Department of Electronics, Information and Bioengineering (DEIB), Politecnico di Milano, Milano, Italy; Department of Information Engineering (DIE), University of Pisa, Pisa, Italy; The BioRobotics Institute and Department of Excellence in Robotics & AI, The BioRobotics Institute, Scuola Superiore Sant’Anna, Pisa, Italy; Department of Brain and Behavioural Sciences, University of Pavia, Pavia, Italy; Digital Neuroscience Centre, IRCCS Mondino Foundation, Pavia, Italy

**Keywords:** Parkinson’s disease, spiking neural networks, basal ganglia, cerebellum, mass models

## Abstract

Parkinson’s disease (PD) is a chronic degenerative disorder of the central nervous system that affects the motor system. The discovery that PD motor symptoms result from the death of dopaminergic cells in the substantia nigra led to focus most of PD research on the basal ganglia. However, recent findings point to an active involvement of the cerebellum in PD. Here, we have developed a multiscale computational model of the rodent brain’s basal ganglia-cerebellar network. Simulations showed that a direct effect of dopamine depletion on the cerebellum must be taken into account to reproduce the alterations of PD neural activity, particularly the increased beta oscillations widely reported in PD patients. Moreover, dopamine depletion indirectly impacted spike-time-dependent plasticity at the parallel fiber-Purkinje cell synapses, degrading associative motor learning as observed in PD. Overall, these results suggest a relevant involvement of cerebellum in PD motor symptoms.

**Significance Statement:** This study highlights the role of cerebellum in Parkinson’s disease (PD). While most studies on PD concentrate on dopaminergic mechanisms in the basal ganglia, here we show that dopamine depletion impacts also on the cerebellum, generating a complex dysfunctional interaction between the two subcortical circuits. To investigate this interaction, we developed *de novo* a multiarea multiscale network model that mechanistically addresses the effects of dopamine depletionon the cerebellum. Our study demonstrates that this aspect is crucial to reproduce experimental data, particularly the increased beta wave activity. Moreover, alterations in spike-time-dependent plasticity at the parallel fibre – Purkinje cell synapse of cerebellum can explain the link between dopamine depletion to motor learning impairment. These simulations indicate that the cerebellum warrants more attention in future PD research.

## Introduction

Parkinson’s disease (PD) is a neurological disorder that results from the degeneration of dopaminergic neurons in the substantia nigra pars compacta (SNc) (1). This reduces dopamine levels in the basal ganglia (BG), modifying local circuit computation, and eventually altering thalamo-cortical activity. As a result, cortical oscillations in the *β* (13 − 30 Hz) band increase, providing a reliable clinical marker of the motor state in Parkinsonian patients (2).

While the role of BG in PDhas been intensely investigated, recently the cerebellum has also been implicated in the disorder’s pathophysiology (3, 4). Cerebellar functions and morphology have been shown to be altered in both human (5) and animal (6) PD studies. However, it is not clear how the loss of SNc dopaminergic neurons might affect cerebellar dynamics and impact the disease.

A recent hypothesis is that the equilibrium of the cerebellar microzones (i.e., functionally defined longitudinal strips of cerebellar cortex (7–9)) might be modified by the dopaminergic loss in PD. This hypothesis is supported by the presence of dopaminergic inputs from SNc and the expression of dopamine receptors in the cerebellum (10). The cerebellum also generates a feedback pathway to dopaminergic nuclei projecting to the ventral caudatus, which explits cerebellar predictive capabilities to regulate the reward system (11, 12). Therefore, the relationship of the cerebellum with the dopaminergic system and the basal ganglia is bidirectional. The cellular apoptosis and reduced activity of the cerebellar cortex in PD (6, 13)) could be instrumental for a the cerebellar role in the disease.

Indeed, there is a di-synaptic connection originating from the subthalamic nucleus (STN) and terminating in the cerebellar cortex via the pontine nucleus (14), thanks to which the synchronous *β*-band oscillations originating from the BG could drive the cerebellum out of its physiological ranges (15–17).

Notably, dopaminergic treatments may enhance cerebellar pathological changes, which are reported to reduce the active dopamine receptors levels in cerebellum (18). Each of these factors may cause modifications of the Parkinsonian cerebellum, and further investigation is needed to understand their relative influence.

Another explanation for the altered cerebellar activity in PD may be attributed to cerebellar compensation to maintain brain activity. Indeed, in early stages of the disorder, patients demonstrated proficient performance in motor tasks despite exhibiting cerebellar hyperactivation in BOLD signal studies (19, 20). Such increase could be explained by a compensation via the cerebello-thalamo-cortical loop for the lower striato-thalamo-cortical circuit activity (21), and these compensatory effects may remain effective until around 70% of the SNc dopaminergic neurons have degenerated (22).

Remarkably, pathological and compensatory effects may represent the causes of different clinical symptoms of PD. Since akinesia and rigidity are rapidly overcome by dopaminergic treatments, they might be induced by the aberrant BG drive on the cerebellum (e.g., (23) showed that levodopa modifies the cerebellum-BG direct pathway). Conversely, tremor might emerge through different mechanisms: the current hypothesis (i.e., the “dimmer-switch” model) formulates that, while BG may trigger the tremor, the cerebellum (specifically the cerebello-thalamo-cortical loop) amplifies tremor amplitude (24). Whether this is caused by compensation or pathological changes in the circuit still needs to be investigated. Unfortunatly,e these findings reflect a relatively recent interest for the cerebellar involvement in PD, leaving the mechanisms behind such altered activity are unclear, hindering the development of more effective pharmacological treatments and neurostimulation treatments (25).

Since it is unpractical to investigate multiple circuit interactions in vivo, the present study anticipates a computational approach to examine the pathological mechanisms responsible for the alterations in cerebellar activity observed in experimental studies. In particular, we modeled the brain regions primarily affected by PD and simulated the electrophysiological hallmarks of the disease.

## Materials and methods

### A computational approach

Brain modeling and simulations are becoming increasingly popular in studying the brain. These tools enable the simulation and study of brain networks and circuits in a controlled environment, offering several advantages over traditional experimental approaches. For instance, computational models allow for the investigation of complex systems that are difficult or impossible to study directly, and they can be used to test hypotheses and generate predictions. Additionally, brain modeling can provide insights into the underlying mechanisms of brain disorders, allowing for the development of targeted interventions and treatments. In this study, a multiarea, multiscale model was designed to simulate the effects of dopamine depletion in spiking neural networks (SNNs). The model included interconnected components, as shown in Figure 1. To achieve this, the study adapted and incorporated spiking models from previous studies for the cerebellar (26) and basal ganglia (27) networks, since these regions were the focus of the investigation. Mass models were also used for the thalamus, thalamic reticular nucleus, and cortex, adapting the work by (28). Spiking neural networks simulate the behavior of single neurons (29), while mass models describe the dynamics of neuronal ensembles (30). By combining these models, we have created a computational tool balancing biological plausibility against the computational burden to examine the interplay between cerebellum and basal ganglia in PD.

**Fig. 1.**
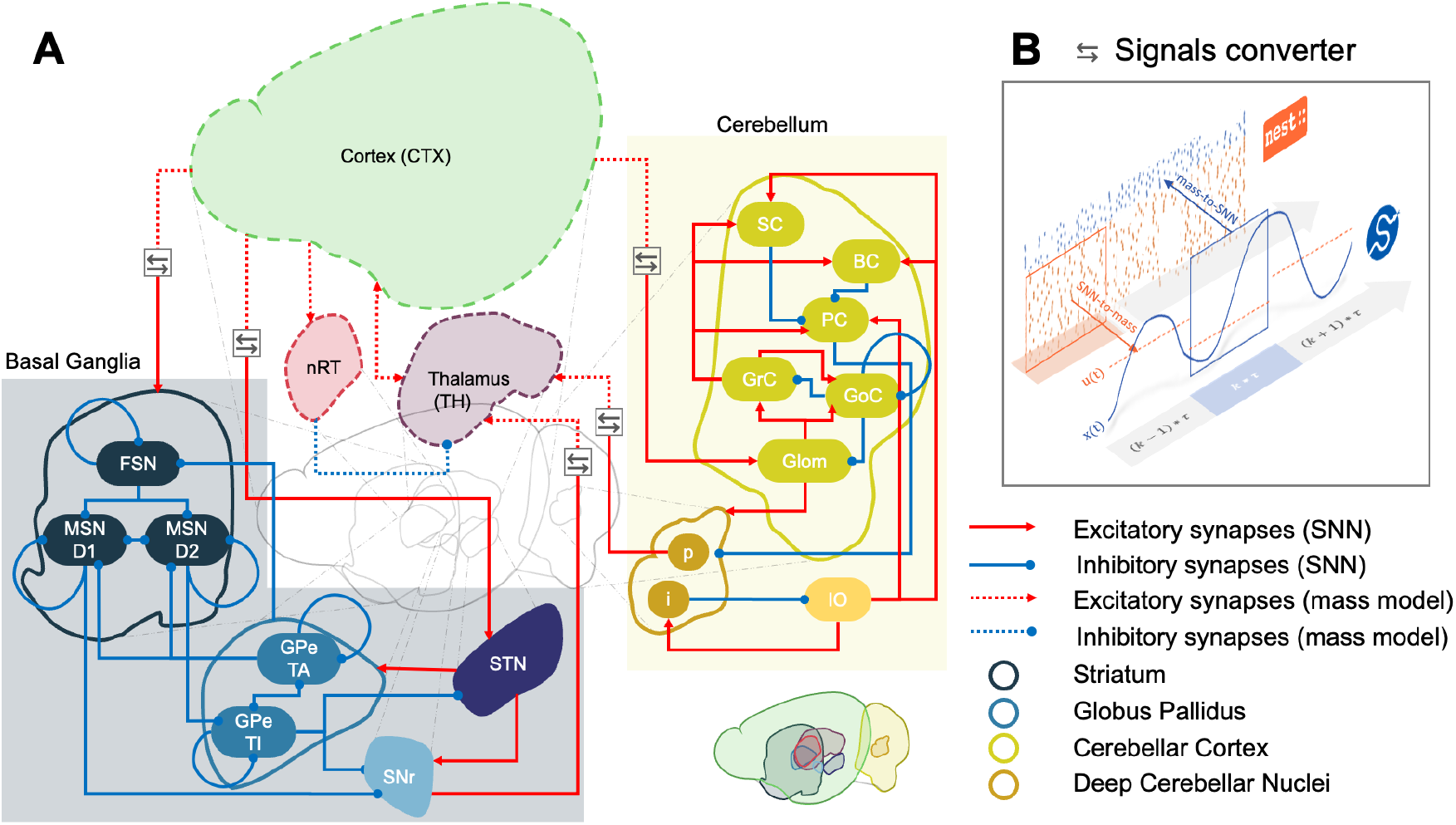
Network architecture. Panel **A** illustrates the connection diagram of the simulated network. The BG (whose subpopulations are represented in different shades of blue) and the cerebellum (shades of yellow) were modeled using spiking neural networks, while the cortex (CTX), the thalamus (TH), and the thalamic reticular nucleus (nRT) were simulated via mass models (indicated by dashed lines). custom software interface (panel **B**) was developed to integrate SNNs and mass models in a multiscale architecture. At each time step of one simulation, output spikes of SNNs are converted to firing rates with a windowed average and given as input to the corresponding mass models. In contrast, mass models’ output firing rates are used as Poisson distribution parameters to compute input spike trains for the connected SNNs. During closed-loop simulations, both the basal ganglia and the cerebellum receive external inputs (not shown in the figure for the sake of clarity) that implicitly mimic the external brain areas’ activity; such inputs target GPe-TI, GPe-TA, SNr, and Glom. List of abbreviations: FSN (Fast Spiking Neurons), MSN-D1 (Medium Spiny Neurons, dopamine 1 receptors), MSN-D2 (Medium Spiny Neurons, dopamine 2 receptors), GPe-TA (external Globus Pallidus, Type arkipallidal), GPe-TI (external Globus Pallidus, Type prototypical), STN (Sub-Thalamic Nuclueus), SNr (Substantia Nigra pars reticulata), Glom (Glomeruli), GrC (Granule Cells), GoC (Golgi Cells), PC (Purkinje Cells), BC (Basket Cells), SC (Stellate Cells), IO (Inferior Olive), p (projecting deep cerebellar nuclei cells), i (inhibitory deep cerebellar nuclei cells).

### Basal ganglia network

The first of the two spiking components of the loop, namely the basal ganglia SNN, was implemented following previous work by (27) that reports detailed neuron and synapse equations. Additionally, in formulating the model, the authors included and validated a parameter to define the dopamine depletion level, making it particularly suitable for this study.

The quantitative computational model of the BG includes the striatal subnetwork, the external Globus Pallidus (GPe), the STN, and the output nucleus, the Substantia Nigra reticulata (SNr). In the striatum, the projection neurons (medium spiny neurons, MSNs) are controlled by the inhibition coming from fast-spiking neurons (FSN) and external Globus Pallidus type A neurons (GPeTA). The model includes dopaminergic neurons in the Substantia Nigra pars compacta (SNc), projecting to striatal neurons and modulating the activity of the entire BG network. Point neuron models were used to ensure a good trade-off between simulation efficacy and biological plausibility: quadratic integrate and fire models with adaptation (31) for FSN and MSN, and adaptive exponential integrate and fire models (32) for the remaining components of the network. Using this formalism, the membrane potential of each neuron in the BG network is described by a set of differential equations that capture the temporal evolution of the voltage guided by the opening and closing of ion channels in the membrane.

Ultimately, this computational model allows the simulation of dopamine depletion mechanisms, whereby the impact of disease progression can be effectively captured by modeling the prominent shifts in both cellular and topological properties of the neural network (for a full description of the dopamine-related changes, see subsection A in Supplementary Information). This is achieved by including a continuous parameter *ξ*, representing the variation of bounded dopamine receptors with respect to the physiological state (*ξ* values range from *ξ* = 0, healthy control, to *ξ* = *−* 0.8, severe PD, with 80% of dopamine receptors unbounded compared to the physiological state).

### Cerebellar network

The second spiking component of the loop, the cerebellum SNN, was adapted from the work by (26). The original model consists of a 3D reconstruction of the olivocerebellar microcircuit, which incorporates optimized neuronal models and morphologically-derived topology (33). This model includes 96,767 neurons and 4,151,182 total synapses, replicating two microzones with their respective olivary nuclei. The model employs Extended-Generalized Leaky Integrate and Fire (EGLIF) point neuron models to simulate neural dynamics and express the main features of cerebellar neurons (34, 35). Although the original 3D extension of the cerebellar microcircuit is not preserved here (our simulation platform is a-dimensional in terms of spatial network organization), the overall structure and connectivity of the network are equivalent to (26).

Such a computationally lightweight model of the cerebellar network includes the three main cerebellar cortical layers as well as the deep cerebellar nuclei (DCN) and the inferior olive neurons (IO). The network receives external inputs via the mossy fibers to glomeruli (Glom) in the granular layer, which connect to granule (GrC) and Golgi (GoC) cells. The subsequent layers include the Purkinje Cells (PC) and molecular interneurons (namely Basket (BC) and Stellate cells (SC)).

PCs receive inputs from the GrCs, both BCs and SCs interneurons, and teaching signals from IO neurons, and in turn, modulate the output of the whole microcircuit through their inhibitory effect on the glutamatergic projecting neurons of the deep cerebellar nuclei (DCNp). The IO neurons are regulated by an inhibitory loop with the GABAergic deep nuclei (DCNi), which control the synchrony and timing of the teaching signal (36).

As the main objective of the present study was investigating the role of the cerebellum in the pathophysiology of PD, we extended the original cerebellar model with a mechanism to account for progressive dopamine depletion based on recent literature findings. Previous studies have indeed suggested that cerebellar cells express dopamine receptors (37), and atrophy of the Purkinje cell layer has been observed in Parkinsonian patients (13). To mimic this effect, we incorporated a simple mechanism of PC apoptosis in the cerebellar model: specifically, we modeled a linear reduction of PC numerosity proportional to the same parameter *ξ* used in the BG SNN (27), with a value of *ξ* = 0 corresponding to the full PC population, *ξ* = *−* 0.8 corresponding to advanced PD. This last case compares well to the 50% PC reduction suggested by a reduction of firing activity in half of the monitored PC following dopaminergic lesion (6).

### Thalamocortical mass models

The remaining areas were modeled using a simplified approach, allowing us to include plausible physiological connections between the detailed cerebellum and the basal ganglia SNNs while maintaining a low computational cost.

For this purpose, we adapted the model composed of seven ordinary differential equations proposed by (28), which describes the temporal dynamics of a cerebellar-basal ganglia thalamocortical network.

The differential equations are formulated based on the Wilson-Cowan approach (30), and can be represented in a compact matrix form (Equation 1)

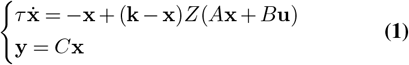

with **x** representing the array of population activities (i.e., number of active neurons at a given time), **u** and **y** the different inputs and outputs of the model, respectively.

In this mathematical formulation, the *n*^*th*^ component of **x** (*x*_*n*_) represents the number of active neurons in the *n*^*th*^ population. Its temporal evolution can be written, following the first equation, as:

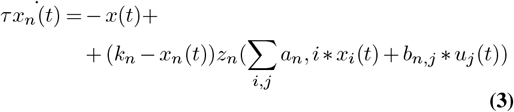

with

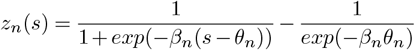

being the response function modeling the portions of cells firing in the *n*^*th*^ population for a given level of input activity *s* (which is the weighted sum of contributions from other *i* populations, Σ*a*_*n,i*_ *x*_*i*_(*t*), and *j* exogenous input sources, Σ _*j*_ *b*_*n,j*_ ∗ *u*_*j*_(*t*)).

The temporal evolution of the *n*^*th*^ population is thus characterized by a set of constants (namely *k*_*n*_, *β*_*n*_, and *θ*_*n*_), which take different values depending on whether the population is excitatory or inhibitory: *k*_*e*_ = 0.9945, *β*_*e*_ = 1.3, *θ*_*e*_ = 4.0 or *k*_*i*_ = 0.9994, *β*_*i*_ = 2.0 and *θ*_*i*_ = 3.7. Finally, *τ* is a time constant and has the same value (10 ms) across all populations. Since both BG and cerebellum are represented by spiking neural network models (hence represented with a more biologically plausible approach), these areas are not included in the system with their own state equations, as in (28), but rather their outputs are fed to the model as exogenous inputs. With these premises, our non-linear time-invariant system resulted in three differential equations (Equation 2), one each for cortex, thalamus, and thalamic reticular nucleus.

The state variables array includes activities from the cortex, thalamus, and thalamic reticular nucleus (**x** = (*x*_*Ct*_ *x*_*T*_ *x*_*n*_) ^*T*^), while the inputs from the spiking models are the action potentials of SNr and DCNp neurons (**u** = (*u*_*SNr*_ *u*_*DCNp*_) ^*T*^),

The weights of the connection are encompassed in sparse matrices where only 4 parameters for *a*_*i,j*_, and 2 for *b*_*i,j*_ are different than zero, accounting for the connectivity among areas in the model.

Furthermore, the cortex is the only area projecting to BG and cerebellum: its activity, *x*_*C*_*t*, modulates the frequency of input spike trains to the *k*^*th*^ spiking population (*y*_*k*_), according to the corresponding connection weight, *c*_*k,Ct*_.

The non-zero weights of the inter-area connections *a*_*i,j*_ were replicated from the original model. At the same time, *b*_*i,j*_ and *c*_*k,Ct*_ were optimized using a genetic algorithm (see Interarea connectivity optimization).

### Multiscale simulation handler tool

Since the mathematical formulation and the software libraries to simulate spiking neural networks and mass models are different (as they operate on distinct scales of resolution), the inputs and outputs of these two classes of models needed to be converted in realtime during simulations to allow neural activity to be transmitted between the different areas in the proposed multiscale network.

To that end, we implemented a custom conversion tool to handle and combine the different signals during simulations, as shown in Figure 1 B: each simulation is iteratively stopped every period *T* to let the SNN and mass models exchange information and set the inputs for the following period, *T* + 1. The instantaneous firing rate from the output SNN population is evaluated as the average number of spikes per second per neuron and becomes the input to the mass model system. On the other hand, the system’s output *y* is used as the instantaneous firing rate of input spike trains to the connected spiking populations, implemented as Poissonian processes with mean *y*.

We based our implementation on a similar dedicated solution which has already been proposed in the “The Virtual Brain” (TVB) framework (38), in the context of managing co-simulation between the TVB, a mass model simulator, and `NEST`, a SNN simulator. The user can set the integration period T, balancing the trade-off between simulation time and results accuracy. We set *T* to 1 and 10 ms in the different cases when we needed, respectively, higher resolution or lower computational load (i.e., the simulation of learning protocols).

### Inter-area connectivity optimization

After defining all the components of the network described above, an optimization process was carried out to fine-tune the weights of the connections between the different areas. The optimization process was carried out through a genetic algorithm specifically designed to minimize the discrepancies between the activity of the multiarea circuit and the results achieved in their corresponding studies, using the following fitness function (Equation 4):

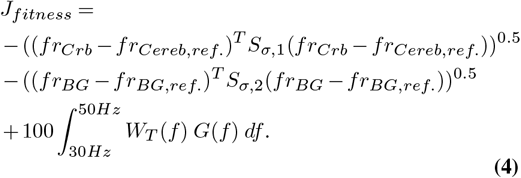

The first two terms of the fitness function are proportional to the Euclidean norm of the difference between the average firing rates of the SNN within the closed-loop model and those observed in stand-alone microcircuits, specifically for the cerebellum (26) and the BG (27) (*fr*_*Crb*_ −*fr*_*Cereb,ref*._ and *fr*_*BG*_ −*fr*_*BG,ref*._), weighted by the inverse of the standard deviations *S*_*σ*_ of the mean firing rates. Since their contribution is considered as a negative term, the more discrepancy the model shows, the lower the fitness function; hence

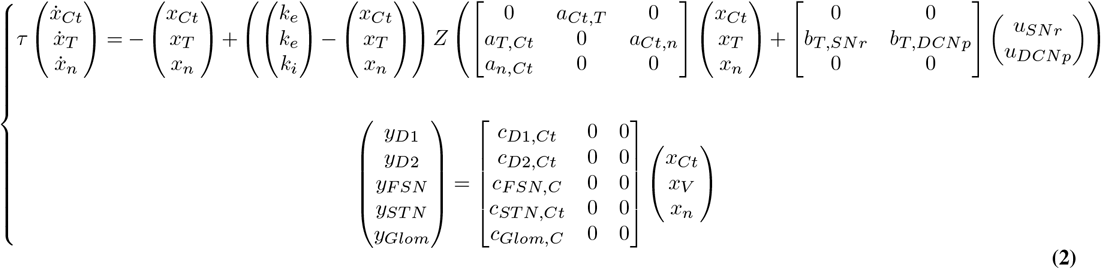

the maximum of *J*_*fitness*_ is found when such differences are closer to zero.

The stand-alone SNNs were first simulated in a state of nonspecific motor activity lasting for 3 seconds: the BG were simulated in an “active” state, as defined in the reference study (27), while the cerebellum was simulated with a Poissonian activity of mean firing rate of 36 Hz given as input, thus using the values reported as conditioned stimulus in the original paper in order to simulate a cerebellar hyperactivation (26).

To avoid overfitting, we relaxed the optimization constraints to only take into account the firing rates from the most informative spiking populations instead of all of them: in particular, the cerebellar populations considered were Glom, DCNp, and PCs, while for the BG, the nuclei taken into account were GPe, STN, and SNr.

Since it was not possible to identify specific reference values for the activity of mass models to be compared with (as the cortical areas have a wide, context-based oscillation regime), the third term of the function expresses the ability of the mass models to reproduce the thalamo-cortical *γ*-band healthy oscillations (*>* 30 Hz) (39), which were also sought to be reproduced in (28). To do so, *J*_*fitness*_ was designed to include the integral of the wavelet transform power of the thalamic *W*_*T*_ activity in the frequencies 30 − 50 Hz, modulated by a Gaussian window *G* (with *mean* = 40 Hz, *SD* = 6 Hz), which increases proportionally to the amount of spectral energy of the signal in the *γ* band.

Once the target values were collected as a reference, the algorithm sought the best combination of connection weights that maximized the fitness function *J*_*fitness*_ by iteratively testing different sets of parameters until a convergence towards the maximum was found (details of the genetic algorithm implementation in Supplementary).

### Testing protocols

To investigate the role of dopamine depletion in the multiarea multiscale model, two testing protocols were conducted in two different conditions of dopamine depletion: simulating dopaminergic effects in the BG only and in both cerebellum and BG.

#### Protocol 1: Active state

We first aimed to examine the effects of dopamine depletion within an active motor state. To achieve this, we repeatedly simulated the multiarea circuit at five levels, from physiological to increasing PD severity (*ξ* = [0, − 0.1, − 0.2, − 0.4, − 0.8]). Specifically, the circuit was simulated 10 times for a period of 3 seconds across the two dopaminergic conditions (only in BG and both cerebellum and BG). The data collected from the simulations were then used to compare the various cases of dopamine depletion across both the temporal and frequency domains by evaluating the variations in the mean firing rates and analyzing the wavelet transforms of the signals from the different areas, and computing the oscillatory phase coherence between the mass models, under all conditions.

#### Protocol 2: Eye Blink Classical Conditioning

The second set of testing protocols aimed at investigating the functional performance of the circuit under both physiological and dopamine-depleted conditions using a motor learning protocol. In fact, we sought to investigate the impact of dopamine depletion on motor learning and plasticity within the cerebellum. To do so, we incorporated a spike-timing-dependent plasticity (STDP) mechanism in the cerebellum SNN in the connection between the parallel fiber and Purkinje Cells (pf-PC) (40) and simulated an eye-blink classical conditioning (EBCC) protocol. The EBCC protocol involves the association of a neutral conditioned stimulus with an unconditioned stimulus, resulting in the acquisition of a learned response to the conditioned stimulus alone (41).

The simulated EBCC experiments consisted of 100 trials of 580 ms, in which the cerebellum SNN received cortical noise at 4 Hz throughout the whole trial, on top of which a complex stimulus at 36 Hz was administered between 100 and 380 ms. Additionally, the cerebellum would receive a teaching signal from the IO at 200 Hz in the 350 − 380 ms window, co-terminating with the conditioned stimulus (CS) (42). The data collected from the repeated trials were analyzed to evaluate learning at the DCNp level in terms of response an-ticipation.

## Data analysis

### Protocol 1: Active state

The behavior during a state of non-specific motor activity was assessed by analyzing both temporal and frequency features. The <u>ma</u>in temporal feature calculated was the mean firing rates (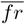) of the spiking models, which summarise populations’ activity over time. According to (43) 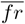 can be calculated as follows:

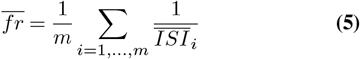

where *m* is the number of neurons in the population and 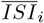 the mean of the inter-spike intervals of the *i*^*th*^ neuron. 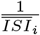 is set to 0 if the *i*^*th*^ neuron spikes less than 2 times during the simulation.

To obtain the frequency features of the signal, First, the signals were converted into the frequency domain using wavelet decomposition (see Supplementary for details), and subsequently the FOOOF (*“Fitting Oscillations and One-Over F”*) technique (44) was used to separate the periodic and aperiodic components of the power spectrum. Using this technique, the aperiodic term can be described by the hyperbolic function 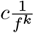: first, the numerical values of the parameters *k* and *c* were estimated using the Python package provided by (44) alongside the paper describing the technique (the `fooof` tool). Then, these components were removed to focus exclusively on the periodic peaks in the spectrum.

Furthermore, we aim to characterize better the emergence of synchronous oscillations in the mass models. To do so, the phase differences between mass models’ activities and the corresponding circular variance values (*CV*) were calculated. The original time series were filtered with a bandpass filter between 13−30 Hz to only keep the beta band components. Then, the phase values *φ*(*t*) of filtered signals were computed from their Hilbert transforms, and pair-wise instantaneous phase differences were calculated for the three mass models in the three dopamine depletion sites (i.e., at time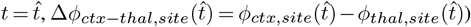.

### Protocol 2: Eye Blink Classical Conditioning

To evaluate the performance of the model during the EBCC, we monitored the activity of two key neuronal populations, Purkinje cells, and deep cerebellar nuclei, as they underwent firing changes driven by plasticity. In particular, we assessed both their intrinsic firing properties, such as firing rate, and their firing modulation, by computing the Spike Density Function (SDF). This method involves convolving individual cells’ spike trains with a kernel function, a Gaussian function of 20 and 10 ms for PCs and DCN, respectively (45), to effectively blur the firing rate estimate over time. This smoothing helps to reduce the effects of random fluctuations in the spike count and provides a more reliable estimate of the underlying firing rate. The SDFs of the neurons within each of the two populations were averaged to obtain the population’s overall SDF.

We also derived the motor output response (the conditioned response, i.e., eyelid closure) by applying a moving average filter with a 100 ms time window to the DCN population SDF to quantify learning further. This was due to take into account the transformation of neural signals into effective eyelid movements. A conditioned response (CR) was detected when the motor output signal reached a fixed threshold of 6.2 Hz in the CR time window and remained above the threshold. The threshold value was selected to have a curve comparable to the one obtained by (42). In this way, each trial corresponds to a boolean variable of success or failure. Lastly, to have a comprehensive measure, we computed the percentage of CRs (%CR) in each block of 10 trials (46, 47).

### Implementation details

The SNN microcircuits were implemented in `Python` 3.8, exploiting the `PyNEST API` to reconstruct and simulate the networks (48). To allow proper integration of the models, both the BG (originally implemented using `NEST Simulator 2.12`) and the cerebellar models (`NEST Simulator 2.18`) were ported to `NEST Simulator 2.20.2`. Furthermore, the mass models’ implementation by (28) was then replicated in `Python` using the `scipy`’s `odeint` routine from its original definition in `MATLAB`. Simulations were performed on a tower computer (64 GB of RAM, Intel(R) Core(TM) i9 CPU 12th Gen) running `Ubuntu OS 20.04.5`.

## Results

### Multiarea circuit activity in physiological conditions

The two spiking networks of the multiarea circuit (cerebellum and BG) had been tuned and validated against *in vivo* data in independent studies (26, 27). Here, we validated the SNNs embedded in the loop using the parameters reported in the original SNN simulations.

*In vivo*, the activity of each brain area is influenced by the activity ongoing in the other areas to which it is (directly or indirectly) connected. Hence, when conducting single-area validations of BG and cerebellum, the impact of other brain areas is indirectly incorporated by how they influence the firing rate of BG and cerebellum networks. In fact, the parameters in the multiarea circuit were adjusted to mimic the firing rates of the BG and cerebellum SNNs, thereby establishing an overarching constraint on the entire multiarea circuit at a global, high-level scale (Figure 2).

**Fig. 2.**
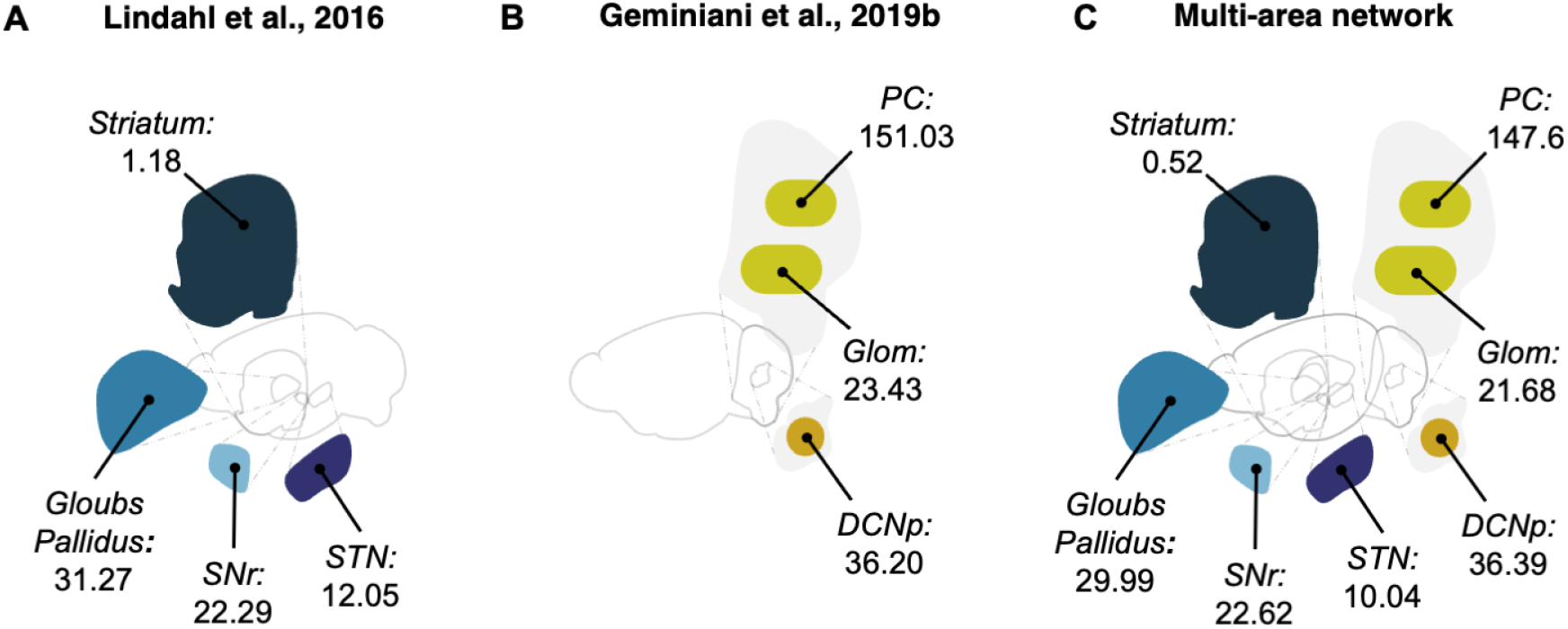
Comparison between firing rates of SNN populations as in stand-alone models and multiarea network. Mean firing rates values (in Hz) of SNN populations in the closed-loop configuration (panel **C**) and when taken individually: BG only in panel **A** (27) and cerebellum only in panel **B** (26).

#### Protocol 1: Active state

To investigate the impact of cerebellar dysfunction in PD on motor processing (in both the temporal and frequency domain), motor learning, and plasticity, we first considered two different pathological hypotheses: dopamine depletion in BG only (27) and dopamine depletion in both cerebellum and BG. The impact of these different sites of dopamine depletion on network activity was compared by evaluating the mean activity of each population in the temporal domain (Figure 3), as well as in the timefrequency domains by analyzing wavelet transforms (Figure 4) and phase coherence.

**Fig. 3.**
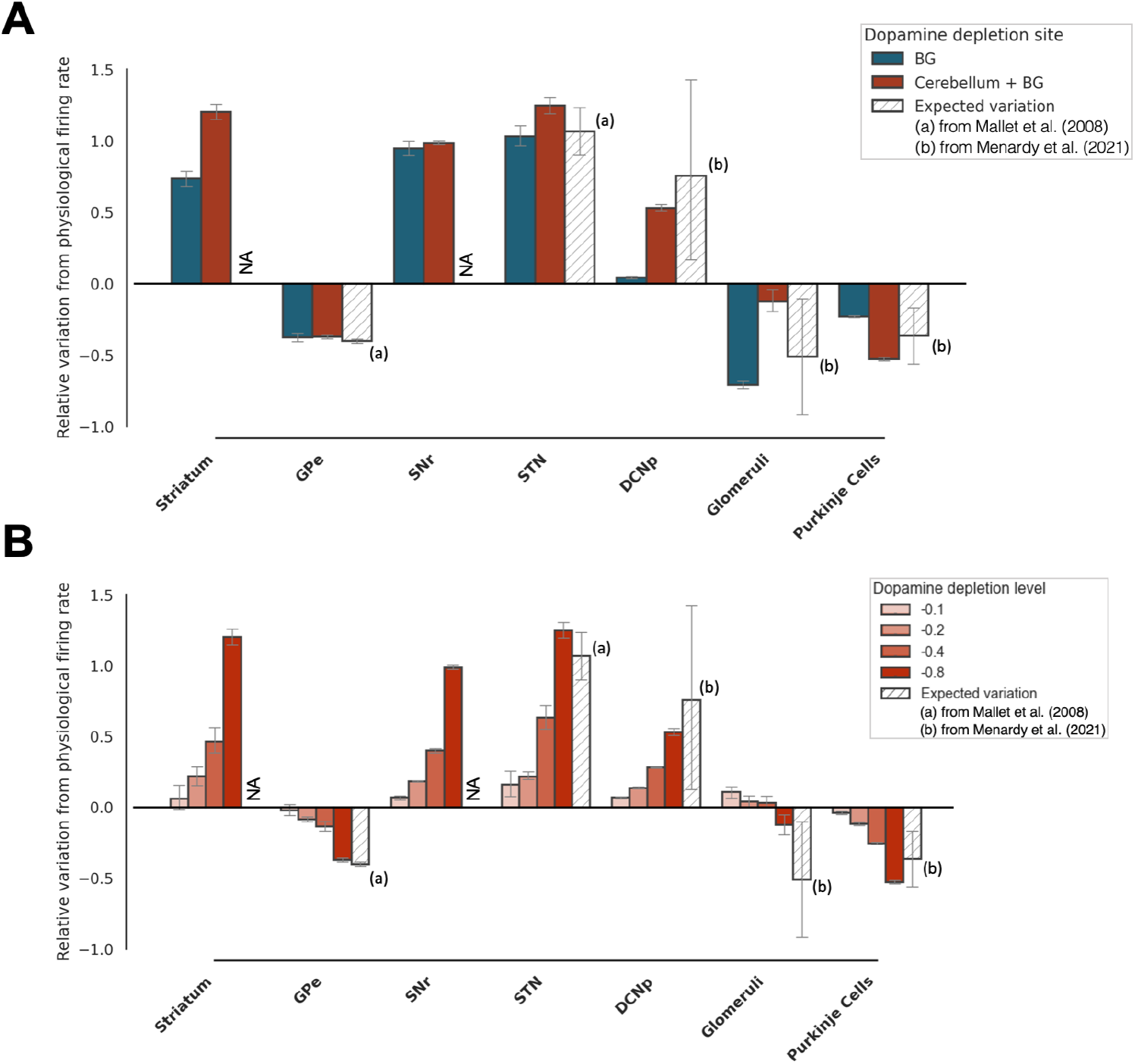
Closed-loop simulations - Firing rate variations with respect to dopamine depletion sites: histograms of normalized firing rate variations in the main spiking populations, grouped by dopamine depletion site and dopamine depletion level. The reported values have been obtained by normalizing the difference between the firing rates in pathological and physiological conditions with respect to the physiological firing rate; therefore, a value of 0 corresponds to no difference between the two, while a value of 1 corresponds to a pathological firing rate that is two times the physiological one. Panel **A** compares the firing rate variation observed with dopamine depletion only in BG (blue) with dopamine depletion in Cerebellum + BG (red); the values are computed considering the most severe pathological scenario (*ξ* = 0.8), as values reported from literature refer to advanced pathology (6, 49). Panel **B** focuses on simulations with dopamine depletion in Cerebellum + BG varying the level of the depletion parameter (*ξ*), i.e., throughout the progression of the pathology.

**Fig. 4.**
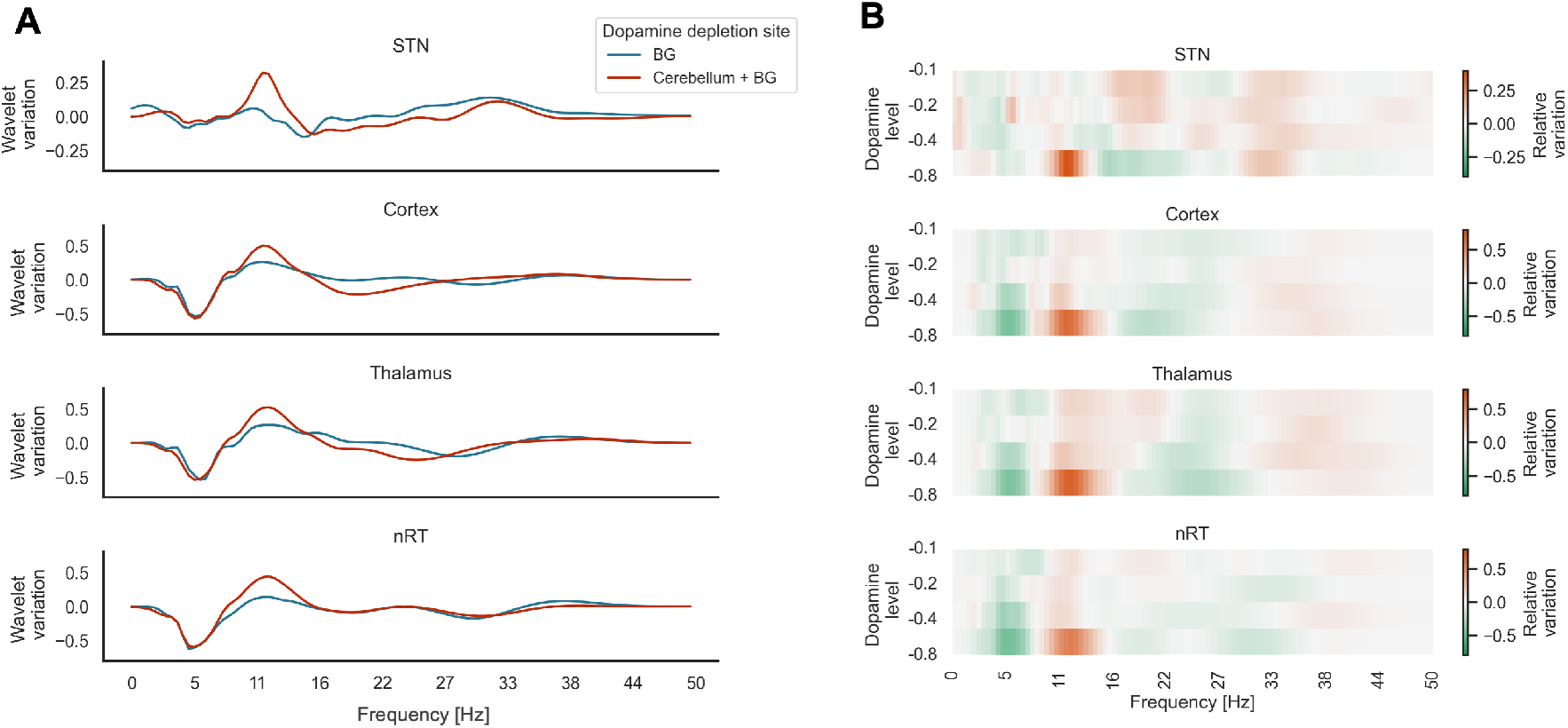
Spectral analysis in different dopamine depletion sites: spectrum values of the three mass models (Cortex, Thalamus, nRT) and subthalamic nucleus. In Panel **A**, each curve represents the variation of the power spectrum under different sites of dopamine depletion (the BG (blue) and both Cerebellum and BG (red)) evaluated at the most severe case (*ξ* = *−*0.8) with respect to the physiological condition 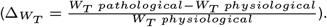 In Panel **B**, the dopamine depletion at both BG and cerebellum is computed at different magnitudes of dopamine depletion (ξ = 0.1, −0.2, −0.4, −0.8).

**Fig. 5.**
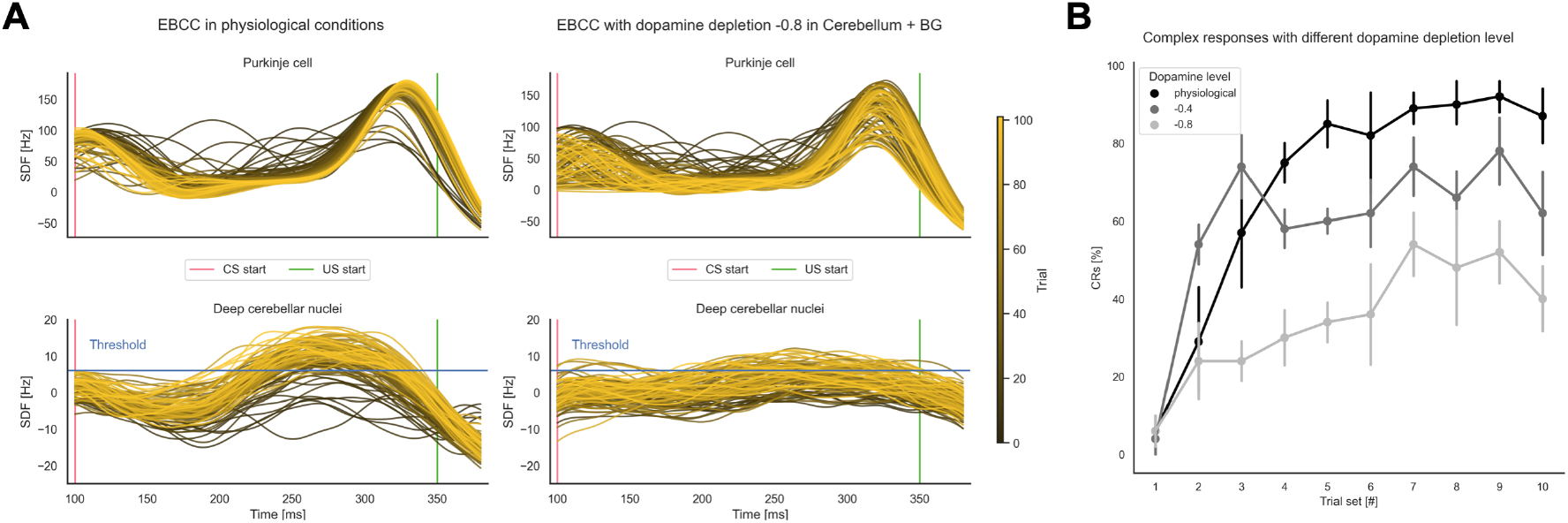
Learning protocol: eye-blink classical conditioning results both in terms of activities of the main populations of interest (PCs and DCNp) and in terms of %CR. Panel **A** shows the results of the EBCC protocol under both physiological (left panel) and dopamine-depleted (right panel) conditions. The entire CS window is shown, with all trials overlaid (each curve represents the SDF within a single trial, and trials’ progression is shown with a shaded color). The signals are adjusted for the baseline evaluated in the 200 ms following the stimuli. Unconditioned stimulus (US) start and threshold for identification of CR are reported. In panel **B**, the percentage of CR identified in each 10-trial block at different magnitudes of dopamine depletion *ξ* (0 = physiological state, −0.4, −0.8)

**Fig. 6.**
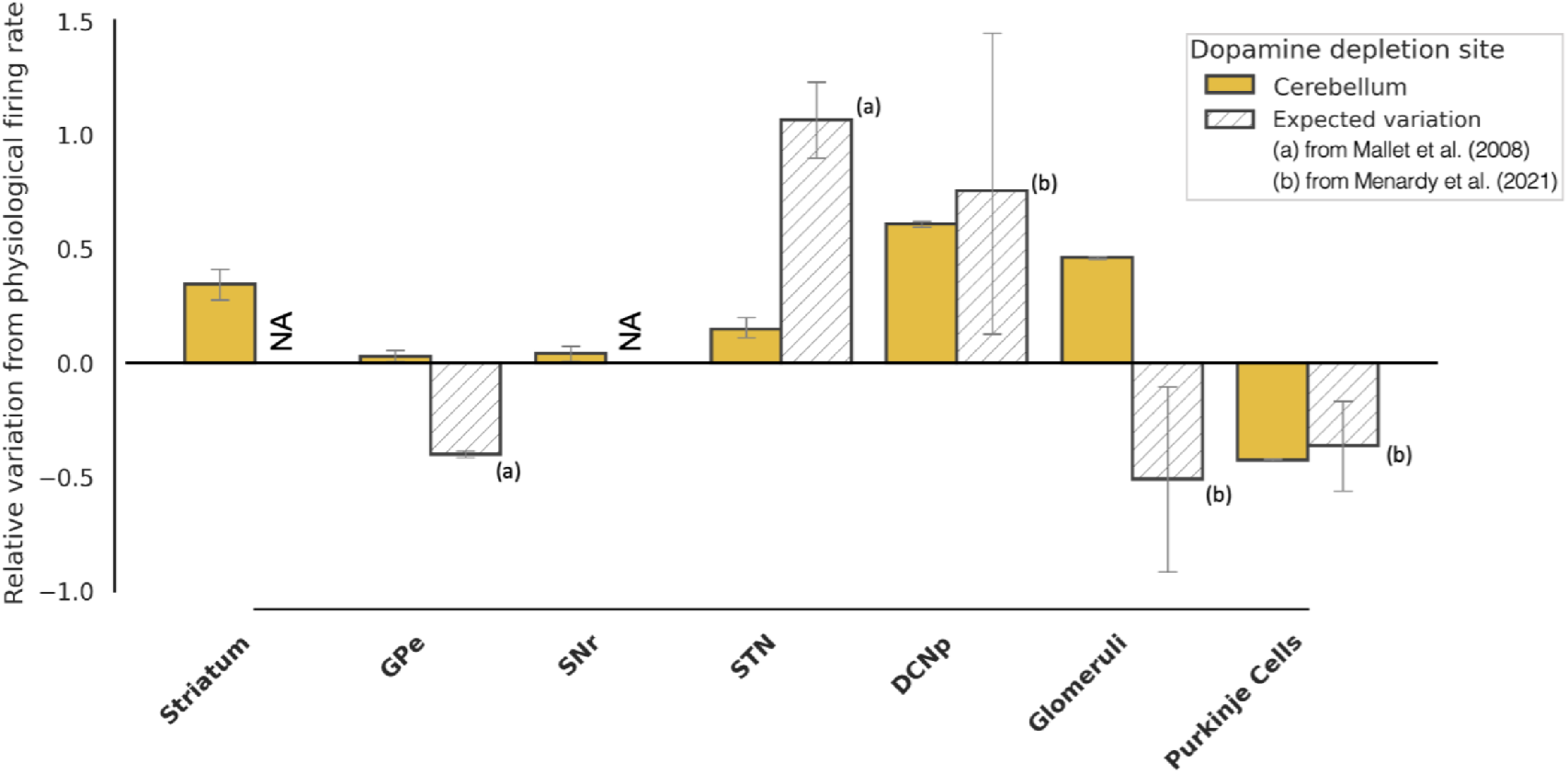
Firing rate variations with dopamine depletion in only Cerebellum: histograms of normalized firing rate variations in the main spiking populations. The reported values are computed considering the most severe pathological scenario (depletion level = 0.8), as values reported from literature refer to advanced pathology (6, 49) and have been obtained by normalizing the difference between the firing rates in pathological and physiological conditions with respect to the physiological firing rate 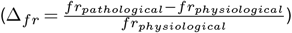 therefore, a value of 0 corresponds to no difference between the two, while a value of 1 corresponds to a pathological firing rate that is two times the physiological one.

The simulation results indicate that dopaminergic effects in BG only are insufficient to explain the experimental firing rate variations in deep cerebellar nuclei (Figure 3 A): the relative difference in mean DCNp activity between reference and simulated values in the BG only condition is 0.763, which decreases to 0.276 in the case of cerebellar and BG condition. It should be noted that the model changes gradually with the intensity of the dopaminergic damage (Pearson correlation *R* ranging from 0.97 to 0.99, *p* − *values* < 0.05 for all SNN populations (N = 20, for each population), with Striatum, SNr STN, DCNp positively correlated and GPe, Glom, and PC negatively correlated). Akin with the concept that the disease is proportional to the severity of the pathological mechanisms (Figure 3 B). Therefore, adding dopaminergic cerebellar mechanisms to Parkinsonian models can improve their descriptive capacities compared to models focusing on the BG alone.

After a thorough temporal analysis of the closed-loop model, we further investigated the underlying neural mechanisms in the frequency domain.

The power spectrum variation with respect to the physiological condition was calculated in the two different conditions of interest, i.e., with dopamine depletion in only BG and in both cerebellum and BG (the maximum depletion value is simulated) (Figure 4 A). We then evaluated the effect of increasing dopamine depletion at both BG and cerebellum (Figure 4 B).

Simultaneous dopamine depletion in the cerebellum and BG led to a most pronounced peak in the beta band, compared to dopamine depletion in BG only: the relative increase, indicated as *median* (*IQR*), of the maximum value in the 13− 30 Hz band from BG only condition to cerebellum and BG condition is 0.33 (0.16) in the Cortex, 0.30 (0.20) in the Thalamus, 0.39 (0.08) in the nRT, 0.40 (0.16) in the STN. Differences between conditions were tested with the Mann-Whitney test and resulted in *p* − *values <* 0.01 for all areas, N = 5 for each population (Figure 4 A). Interestingly, dopamine depletion in the cerebellum only caused a decrease in power spectrum across all frequency values in most areas, except for the STN (Figure 8 in Supplementary).

**Fig. 7.**
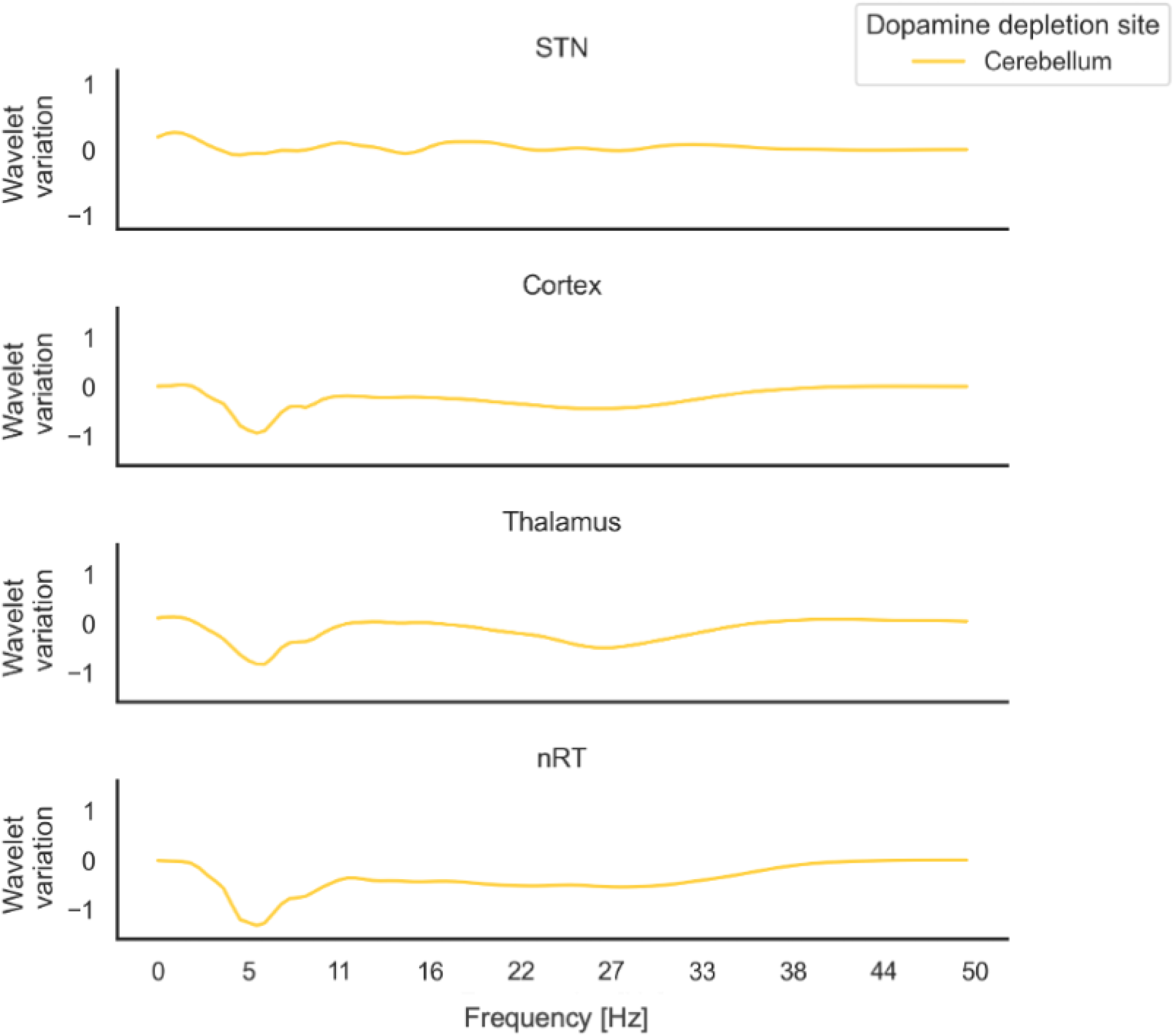
Spectral variation with dopamine depletion in Cerebellum: spectrum values of the three mass models (Cortex, Thalamus, nRT) and subthalamic nucleus (STN). Each curve represents the variation of the power spectrum evaluated at the most severe case (depletion level -0.8) with respect to the physiological condition 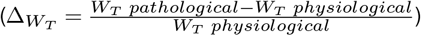.

**Fig. 8.**
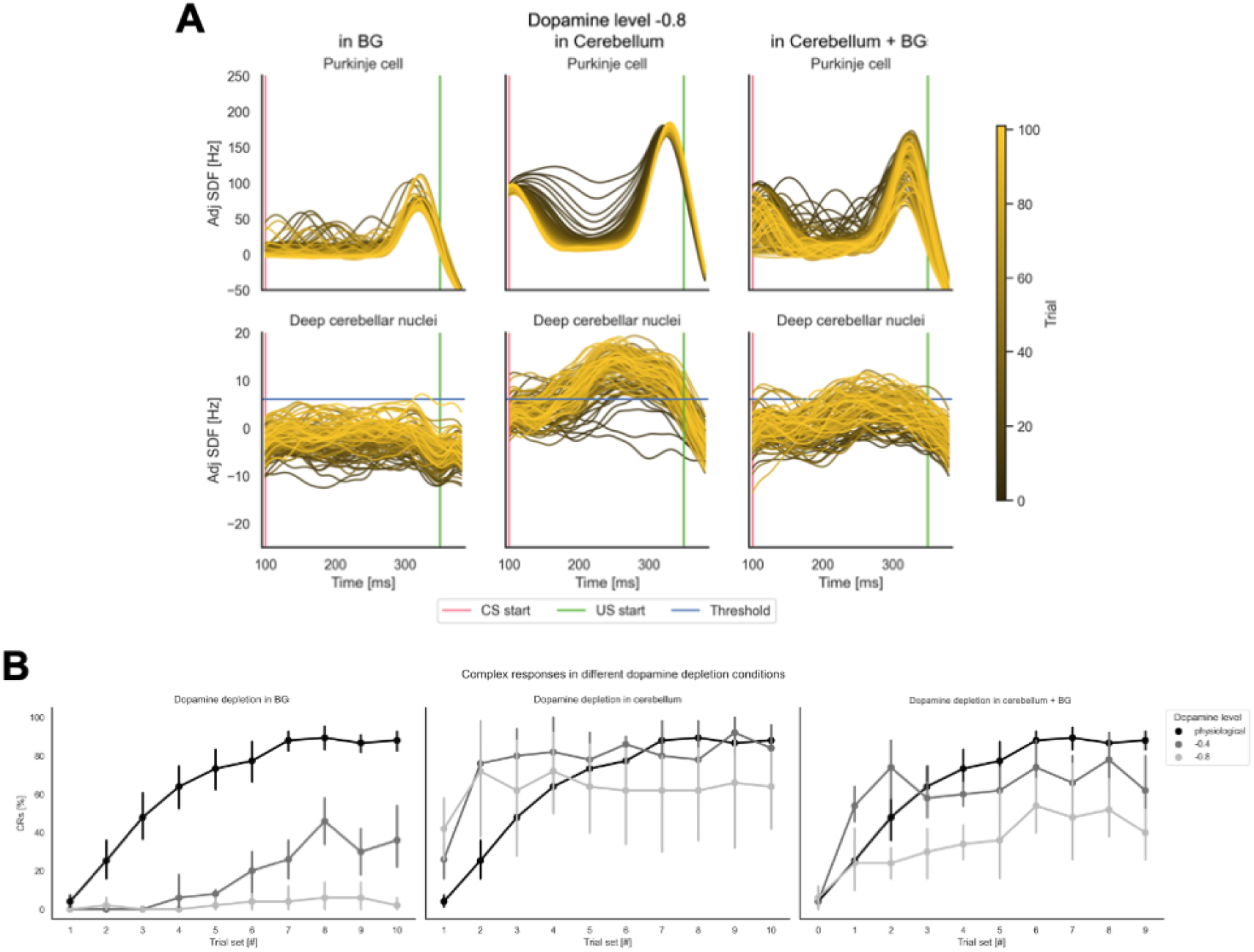
Learning protocol: eye-blink classical conditioning results both in terms of activities of the main populations of interest (PCs and DCNp) and in terms of %CR. Panel **A** shows the results of the EBCC protocol when dopamine depletion is modeled in different sites (only BG on the left, only Cerebellum in the center, Cerebellum + BG on the right) at the most severe case (depletion level -0.8). The entire CS window is shown, with all trials overlaid (each curve represents the SDF within a single trial, and trials’ progression is shown with a shaded color). The signals are adjusted for the baseline evaluated in the 200 ms following the stimuli. Unconditioned stimulus (US) start and threshold for identification of CR are reported. In panel **B**, the percentage of CR identified in each 10-trial block, at different sites (as in panel **A**) and magnitudes of dopamine depletion (0 = physiological state, -0.4, -0.8) are overlaid.

We analyzed the circular variance to further elucidate the emerging synchrony between time-varying signals in the *β* band. Simulations revealed that stronger corticothalamic synchronization (i.e., a higher Circular Variance) was achieved when dopaminergic effects were modeled in both the cerebellum and BG (*CV*_*BG*_ = 0.191, *CV*_*cerebellum*+*BG*_ = 0.132, see Figure 9 in Supplementary).

**Fig. 9.**
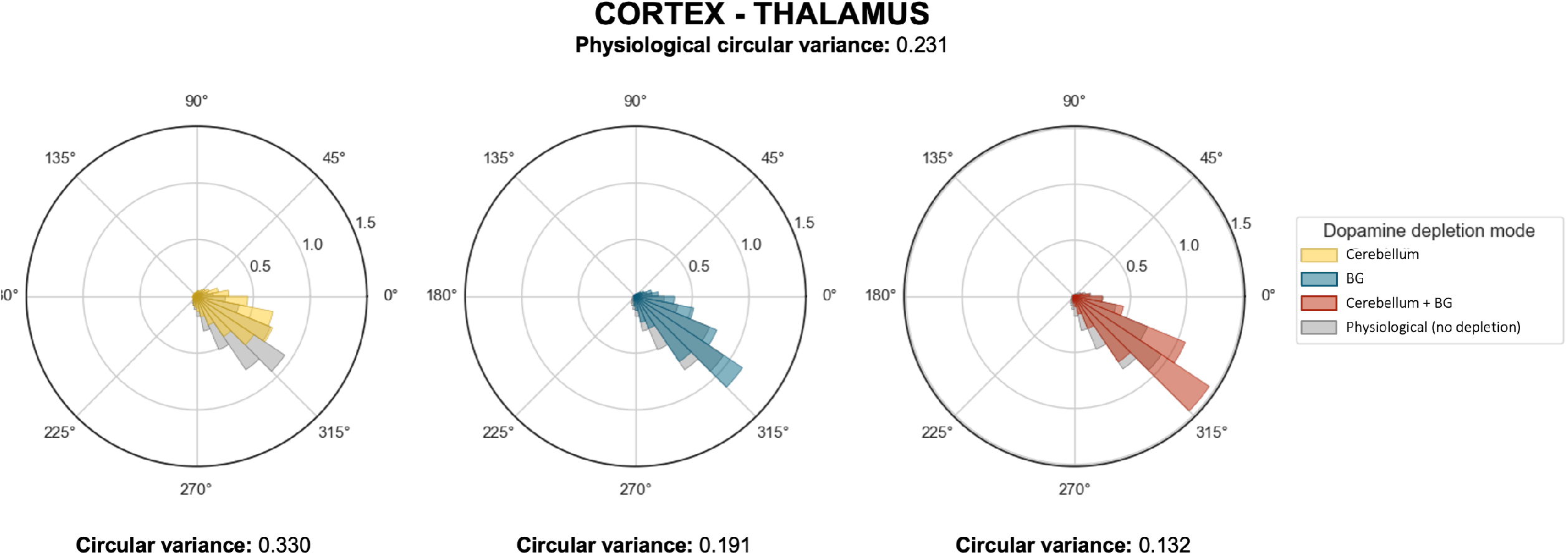
Circular variance of mass models’ activities: polar histograms of the phase differences between mass models’ activities and the corresponding circular variance values in three different sites of dopamine depletion (Cerebellum, BG, and Cerebellum + BG, with depletion level *ξ* = −0.8 and in physiological conditions).

#### Protocol 2: EBCC

Eyeblink classical conditioning (EBCC) was simulated to investigate the effects of dopamine depletion on motor learning and plasticity in the cerebellum by incorporating a mechanism for STDP in the pf-PC connection (40).

According to Figure 5, we focused the EBCC protocol on the model with the highest impact of dopamine effects, i.e., with depletion affecting both BG and cerebellum. Simulations were performed with different degrees of dopamine depletion, comparing the motor learning performance to that obtained under physiological conditions. The same analysis results obtained using dopamine depletion in BG only or in cerebellum only are presented in Supplementary.

In physiological conditions (on the left in Figure 5 A), we observed an initial potentiation coinciding with the start of the CS, followed by a gradual depression towards the end of the CS (due to long-term depression) that anticipates the onset of the US, corresponding to a decrease of 36.8 (3.59) Hz (N = 5) between the averaged activity of the first 10 trials and the last 10 trials in the 280 − 380 ms window. This behavior is reflected in the DCNps, which are progressively released from PCs inhibition along the trials, anticipating the response and exceeding the threshold that activates the CR.

In the PD case (on the right in Figure 5 A), two main effects on PCs and DCNps were observed. On one hand, the PC input showed higher variability, and the learning was less marked, corresponding to a decrease of 29.9 (4.45) Hz (N = 5). This was caused by the reduced drive caused by the dopamine depletion at the BG level and reflected in a reduced cortical input to the cerebellum and by dopamine depletion in cerebellum that reduced the number of healthy PCs. On the other hand, the output from the DCNps was reduced because of the reduced drive from the cortex via the mossy fibers. Finally, we quantified the conditioned response rate, i.e., the performance of the cerebellum in anticipating the response, in both physiological and dopamine-depleted conditions (*ξ* =− 0.4 and *ξ* = − 0.8) (Figure 5 B). As we compared the results to the physiological case, which is in line with experimental values (47), we found that the addition of dopamine depletion led to a decline in performance, which worsened as the depletion progressed (%CR decrease at the last trial set of 30 (20)% from physiological to mild PD states and of 40 (20)% to severe PD states, N=5).

## Discussion

This study sets the computational basis of abnormal multiarea interactions in PD bringing the cerebellum (3, 4, 6) to the forefront.

Incorporating the cerebellum into the closed-loop multiarea circuit provides compelling evidence showcasing how dopamine-dependent cerebellar dysfunction intensifies beta oscillations and hampers motor learning in PD. This finding highlights the need to extend the pathophysiology of PD symptoms to multiple brain regions beyond the basal ganglia, both in experimental and computational models, with the cerebellum acting as a key player.

The multiscale approach adopted here required the integration of multiple spiking neural networks and neural masses. The multiarea model passed a rigorous validation process. Indeed, it successfully replicated the activity of existing cerebellar and BG models (26, 27), which were previously studied in isolation and validated against *in vivo* data. We deem this validation of the highest relevance to reliably capture the interactions among multiple brain regions acting in closedloop, generating *de facto* an innovative computational approach to the investigation of multiple brain circuit interactions (5, 14).

By challenging the multiarea model in both physiological and Parkinsonian conditions, we explored the physiology of the cerebellum -basal ganglia interactions (19, 20). Following incorporation of dopamine-dependent dysfunction in the cerebellum (13), as well as in the basal ganglia (27), the simulations yielded results consistent with experimental observations (6, 49), providing compelling evidence for a direct cerebellum’s involvement in PD-related dynamics. The internal degeneration of the cerebellar cortex (i.e., PC apoptosis) impacted the beta oscillations uncovering a pronounced low-frequency peak in the beta band. These simulations thus identify the cerebellum as a core mechanisms for the reinforcement of aberrant neural activity through the cerebellothalamo-cortical loop in PD (24).

Taking our investigation a step further, we delved into functional motor learning by simulating a virtual eyeblink classical conditioning task (41). Remarkably, despite the altered cellular structure, the cerebellum demonstrated an adaptive capacity, albeit with reduced efficacy compared to the physiological state. This finding highlights the cerebellum’s ability to partially compensate for dopamine-dependent changes (20) and underscores its importance in motor learning and adaptation processes (50).

### Comparison with similar models

It is worth noting that the proposed implementation does not represent the only multiarea, multiscale model investigating PD or the interactions between BG and cerebellum during motor learning.

A study from (51) developed a multiscale model comprising spiking BG and thalamo-cortical mean-field areas implemented using the TVB framework (38) to investigate the network effects of deep brain stimulation (DBS) on the STN. The resulting model generates biologically plausible activity in resting state and during virtual DBS, resulting in a promising approach that could be developed as a clinical platform for neurosurgeons and used to design therapies for individual patients before surgical interventions.

Another interesting proposal has been recently advanced by (52), who developed a systems-level computational model of motor learning, including a cortex-basal ganglia motor loop, the cerebellum, and central pattern generators representing the brainstem. The model can learn arm movements toward different motor goals and reproduce human data in a motor adaptation task with cognitive control, sharing our findings regarding the fundamental importance of building models that account for the interplay between BG and cerebellum. Although both studies represent essential milestones towards a better understanding of both PD and motor learning, they differ from our model in some crucial aspects: although the network in (51) is composed of multiple cortical nodes leveraging the “TVB” framework, however, it does not model the cerebellar contribution explicitly, nor investigates functional motor learning impairments in PD. (52), on the other hand, provide a precise functional interpretation of the BG-cerebellum interplay and their model can produce complex motor responses that are close to human motor adaptation data. Still, their system-level design abstracts remarkably from the brain regions it represents and the authors do not consider any pathological scenario. With this being said, integrating the salient aspects of these two models with our network in a holistic representation would surely improve the descriptive (and possibly predictive) power of the proposed model.

### Limitations and future developments

Although the results are satisfactory, the proposed model has some limitations that shall be addressed with further refinements aimed at improving its biological plausibility. First, not all the interactions between BG and cerebellum have been modeled. Specifically, the cerebellum sends di-synaptic inputs to the putamen through the thalamus, while outputs from the STN reach the cerebellar cortex through the pontine nuclei. However, there are no cues about the functional role of these connections (14), so their use in the models would require extensive research and careful hypotheses.

Furthermore, no dopaminergic effects other than the PCs apoptosis were considered when simulating the functional motor task, and only the cerebellum was equipped with synaptic plasticity. Thus, in order to fully untangle the interplay between the BG and the cerebellum in motor processing, the dopamine-based reward signals in both the BG (that are essential for their reinforcement learning properties) (53) and in the cerebellum (given the presence of dopamine receptors in the region (10, 54)) should be introduced in the spiking model and varied according to the severity of the simulated Parkinsonian conditions.

The thalamocortical areas have been modeled using a highly simplified approach reducing the computational cost. Deploying the present model on dedicated hardware would allow converting these structures into detailed spiking models. These, in turn, could be used to investigate further the cerebellar impact on the oscillatory drive exerted by BG on the thalamo-cortical circuit in PD patients. In light of this, the present simulator could be adapted to run in the TVB framework (38).

Lastly, since the outcome of the simulations could only be partially compared to experimental findings due to lack of data, the model is highly predictive in nature: in order to fully validate it, specific experiments shall be conducted to compare the model outcomes to signals obtained *in vivo*, although acquiring robust neural activity data from the same subject during the progression of the disease is a challenging task.

To conclude, our model highlights the importance of considering the cerebellum in experimental and computational studies aimed at investigating PD and represents an improvement over previous models that only reproduced a single area (27, 55), using oversimplified mass models (28) and did not explicitly model the cerebellum (56). Our approach paves the way for a unified physiopathological framework explaining Parkinsonian brain activity quantitatively across complexity scales.

## Code Availability

The code used for this study can be freely accessed via the following GitHub repository: https://github.com/benedettagambosi/PD_BG_Cereb_loop.

## ACKNOWLEDGEMENTS

This project has received funding from the European Union’s Horizon 2020 Framework Program for Research and Innovation under the Specific Grant Agreement No. 945539 (Human Brain Project SGA3). AA is funded by the Project “EBRAINS-Italy (European Brain ReseArch INfrastructureS-Italy),” granted by European Union – NextGenerationEU (Italian PNRR, Mission 4, “Education and Research” - Component 2, “From research to Business” Investment 3.1 - Call for tender No. 3264 of Dec 28, 2021, of Italian Ministry of University and Research), Project code IR0000011, Concession Decree No. 117 of June 21, 2022, adopted by the Italian Ministry of University and Research, CUP B51E22000150006).

## Supplementary Information

### A. Dopamine depletion

The effects of dopamine depletion in the BG network model have been replicated from the original study (27). A continuous parameter *ξ* ranging from 0 (physiological dopamine level) to − 0.8 (low dopamine level) is included in the parameters, controlling some characteristic changes in network activity related to dopaminergic effects. Moreover, a scaling coefficient for the corresponding parameter fitted from experiments, which determines the relationship between dopamine receptor occupancy and the magnitude of the effect, is included (Table 8).

**Table 1:**
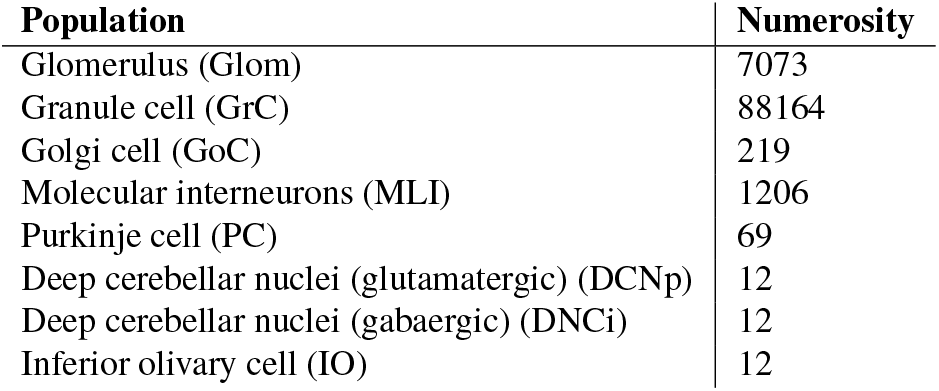
Cerebellar Numerosity.

**Table 2:**
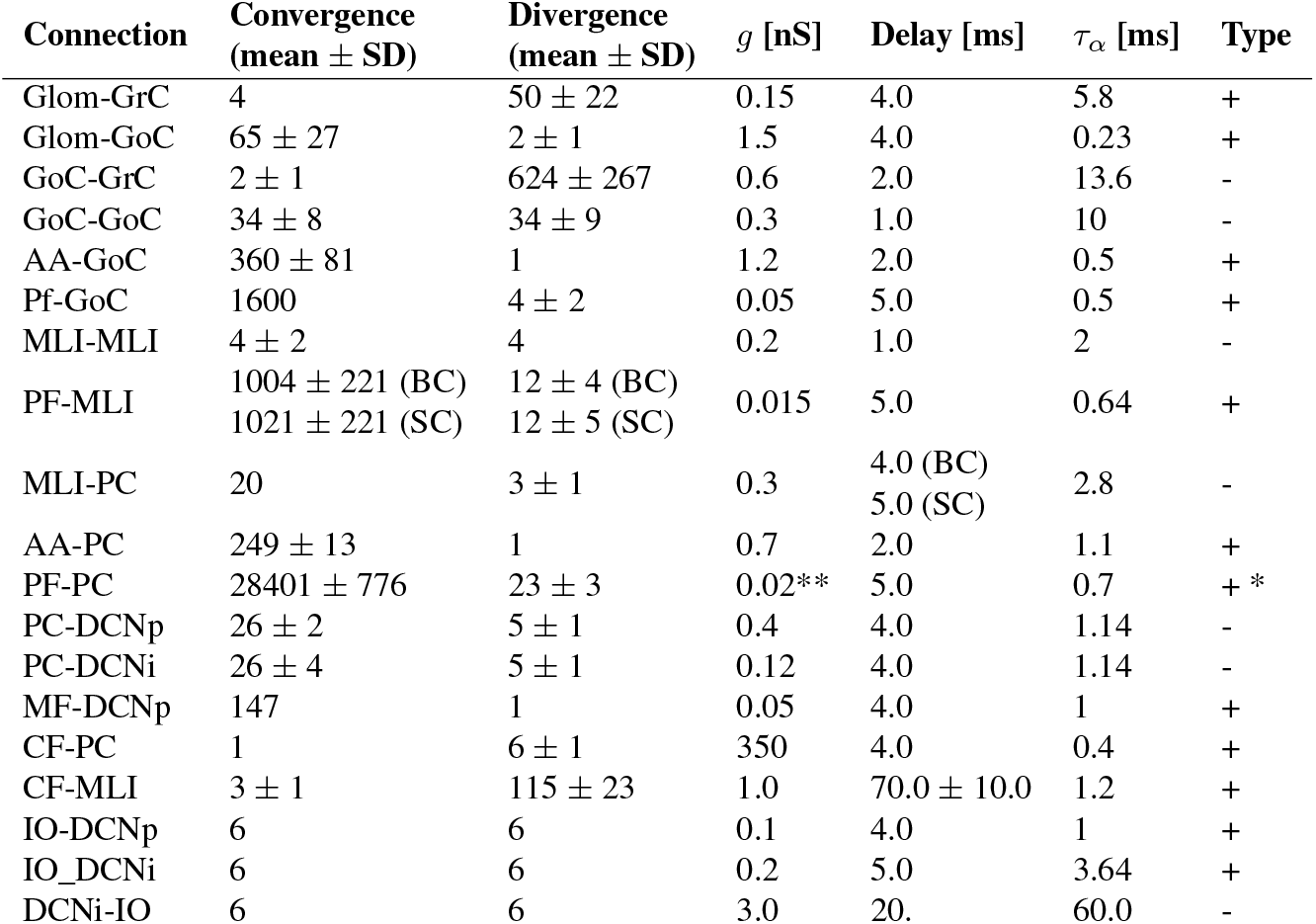
**Cerebellar connections parameters**: *τ*_*α*_ = time constants of the conductance, *g* = conductance. Adapted from (26). Type is + for excitatory and for inhibitory connections. **\*** plastic synapse defined as in (40). ** adapted value to reproduce learning.

| Connection | Convergence<br>(mean ± SD) | Divergence<br>(mean ± SD) | $g$ [nS] | Delay [ms] | $\tau_\alpha$ [ms] | Type |
| --- | --- | --- | --- | --- | --- | --- |
| Glom-GrC | 4 | 50 ± 22 | 0.15 | 4.0 | 5.8 | + |
| Glom-GoC | 65 ± 27 | 2 ± 1 | 1.5 | 4.0 | 0.23 | + |
| GoC-GrC | 2 ± 1 | 624 ± 267 | 0.6 | 2.0 | 13.6 | - |
| GoC-GoC | 34 ± 8 | 34 ± 9 | 0.3 | 1.0 | 10 | - |
| AA-GoC | 360 ± 81 | 1 | 1.2 | 2.0 | 0.5 | + |
| Pf-GoC | 1600 | 4 ± 2 | 0.05 | 5.0 | 0.5 | + |
| MLI-MLI | 4 ± 2 | 4 | 0.2 | 1.0 | 2 | - |
| PF-MLI | 1004 ± 221 (BC) | 12 ± 4 (BC) | 0.015 | 5.0 | 0.64 | + |
| | 1021 ± 221 (SC) | 12 ± 5 (SC) | | | | |
| MLI-PC | 20 | 3 ± 1 | 0.3 | 4.0 (BC) | 2.8 | - |
|  |  |  |  | 5.0 (SC) |  |  |
| AA-PC | 249 ± 13 | 1 | 0.7 | 2.0 | 1.1 | + |
| PF-PC | 28401 ± 776 | 23 ± 3 | 0.02** | 5.0 | 0.7 | + * |
| PC-DCNp | 26 ± 2 | 5 ± 1 | 0.4 | 4.0 | 1.14 | - |
| PC-DCNi | 26 ± 4 | 5 ± 1 | 0.12 | 4.0 | 1.14 | - |
| MF-DCNp | 147 | 1 | 0.05 | 4.0 | 1 | + |
| CF-PC | 1 | 6 ± 1 | 350 | 4.0 | 0.4 | + |
| CF-MLI | 3 ± 1 | 115 ± 23 | 1.0 | 70.0 ± 10.0 | 1.2 | + |
| IO-DCNp | 6 | 6 | 0.1 | 4.0 | 1 | + |
| IO_DCNi | 6 | 6 | 0.2 | 5.0 | 3.64 | + |
| DCNi-IO | 6 | 6 | 3.0 | 20. | 60.0 | - |

**Table 3:**
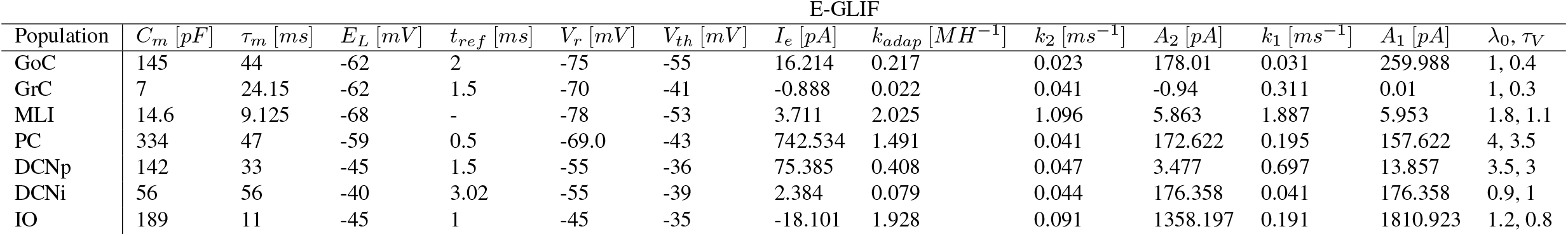
**Extended generalized leaky integrate and fire parameters**: *C*_*m*_ = membrane capacitance; *τ*_*m*_ = membrane time constant; *E*_*L*_ = resting potential; *t*_*ref*_ = refractory period; *V*_*r*_ = reset potential; *V*_*th*_ = threshold potential; *I*_*e*_ = endogenous current; *k*_*adap*_, *k*_2_ = adaptation constants; *k*_1_ = decay rate of the intrinsic depolarizing current; *A*_2_, *A*_1_ = model currents update constants; *l*_0_, *τ*_*V*_ = escape rate parameters. As in (34).

**Table 4:**
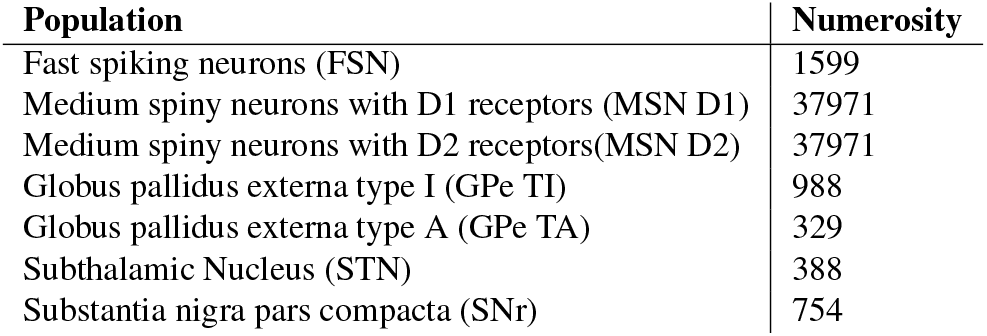
Basal Ganglia Numerosity.

**Table 5:**
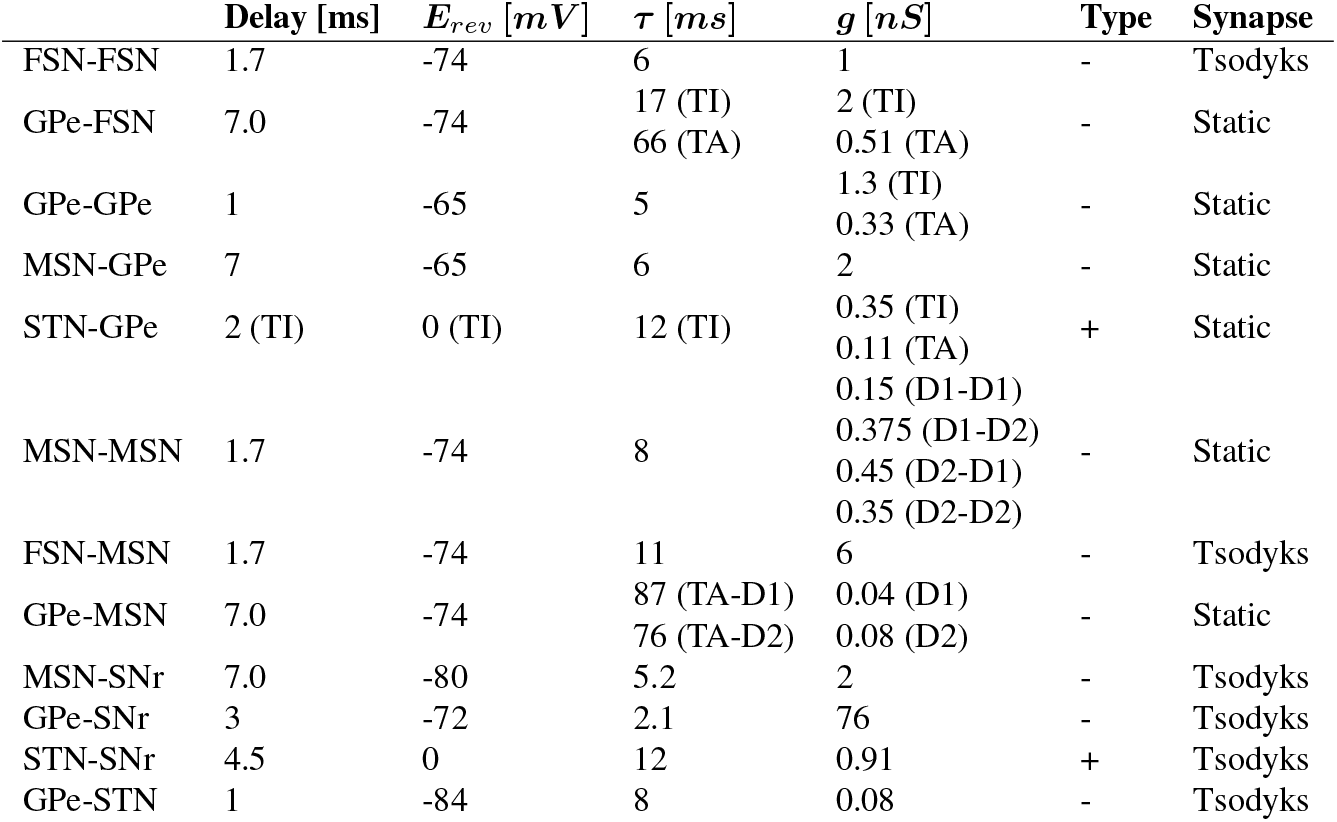
**BG connections parameters**: *E*_*rev*_ = reversal potential; *τ* = time constants of the conductance, *g* = conductance. As in (27).

**Table 6:**
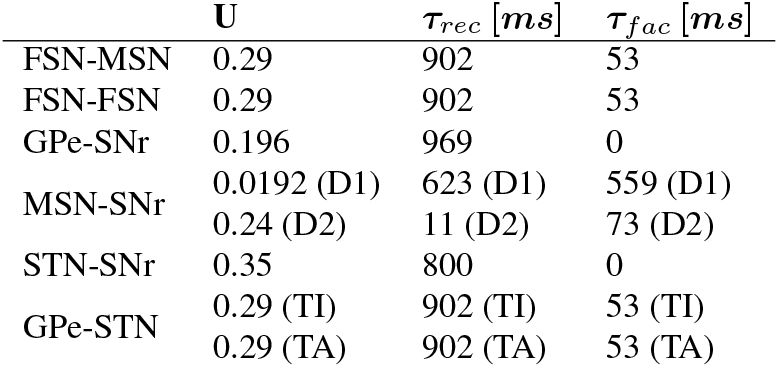
**BG Tsodyks synapse parameters**: *U* = utilization factor ; *τ*_*rec*_ = recovery time constant, *τ*_*f ac*_ = facilitation time constant. As in (27).

**Table 7:**
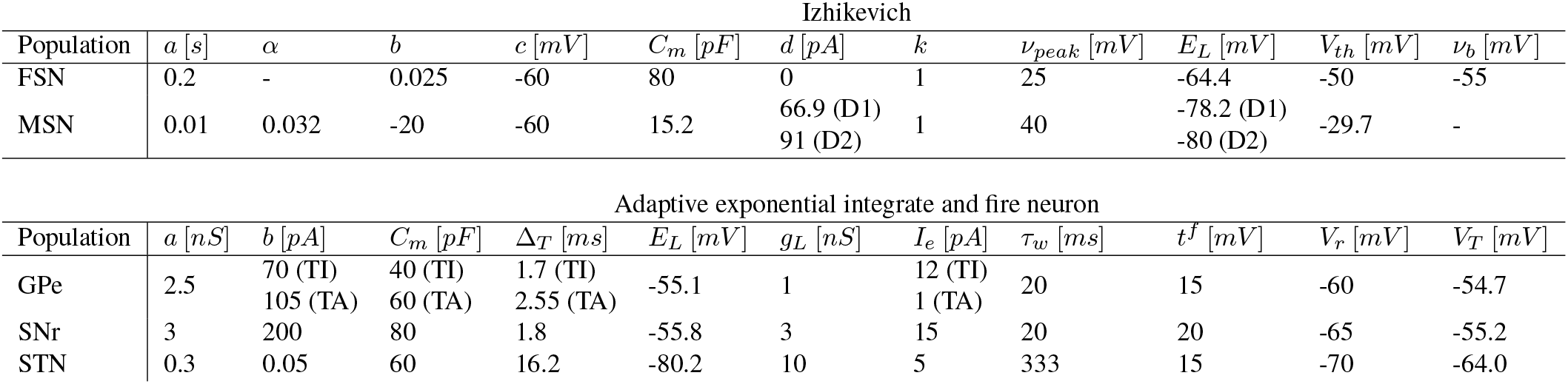
**Izhikevich neuron parameters** *a* = Recovery current time constant; *b* = Spike-triggered adaptation; 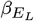 = Magnitude of D1 effect on resting potential; *C*_*m*_ = membrane capacitance; Δ_*T*_ = Slope factor of spike upstroke; *E*_*L*_ = Resting potential; *g*_*L*_ = Leak conductance; *I*_*e*_ = current injected to have desired fr (8*Hz* for TA, 18*Hz* for TI); *t*^*f*^ = Spike cutoff; *V*_*r*_ = reset potential. **Adaptive exponential integrate and fire neuron parameters** *a* = Subthreshold adaptation; *b* = Voltage dependency of recovery current; *c* = Spike reset; *C*_*m*_ = membrane capacitance; *d* = Summed recovery current contribution following an action potential; *k* = Steady-state voltage dependence; *ν*_*b*_ = Voltage dependence recovery current; *ν*_*peak*_ = Spike cutoff; *v*_*r*_ = Resting potential; *V*_*th*_ = Threshold potential; *τ*_*w*_ = Adaptation time constant. As in (27)

**Table 8:**
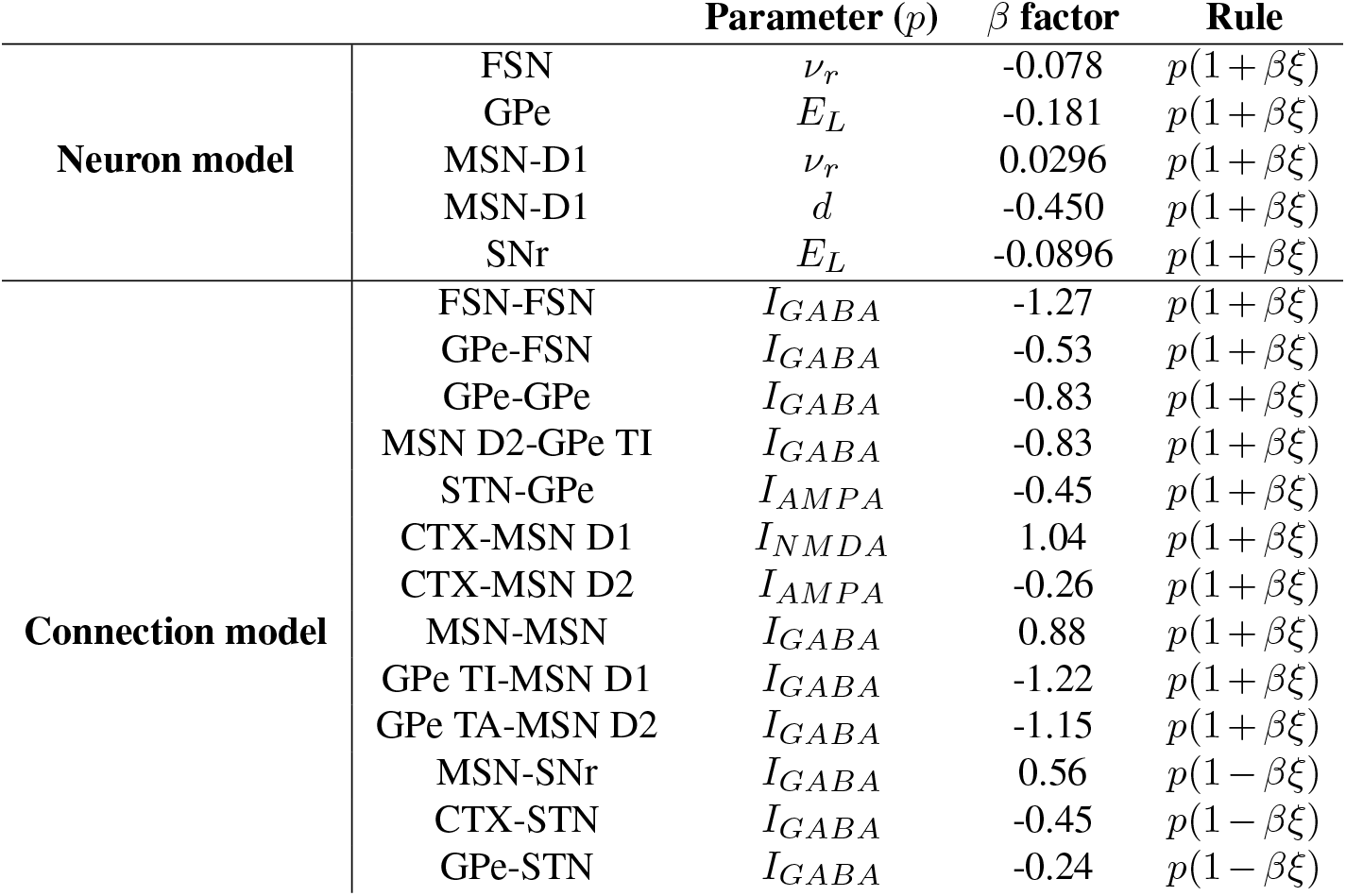
**Multiplicative factor controlling effects of dopamine depletion**: *v*_*r*_, *E*_*L*_ = resting potential; *d* = Summed recovery current contribution following an action potential; *I*_*GABA*_, *I*_*NMDA*_, *I*_*AMPA*_ = GABA, NMDA, AMPA currents (defined as *I* = *g*(*E*_*rev*_ *− V*), with *g, E*_*rev*_ and *V* being the conductance, the reversal potential and the membrane potential correspondingly)

### B. Genetic algorithm

The genetic algorithm was implemented with the PyGAD library (57), with the following settings

- number of generations (i.e., the number of iterations that the genetic algorithm will run for): 5;
- number of parents (i.e., the number of solutions that will be selected for mating and producing offspring in each generation): 2;
- population size (i.e., the number of candidate solutions that will be evaluated in each generation): 2;
- number of genes (i.e., the number of parameters that make up each candidate solution): 4;
- parent selection method: steady-state selection;
- mutation type: random (i.e., each gene in a candidate solution has a probability of being randomly mutated);
- genes to mutate: 50 % ;
- crossover: single point;
- no stopping criteria.

