## supplementary file for "Dopamine-dependent cerebellar dysfunction enhances beta oscillations and disrupts motor learning in a multiarea model"

### Supplementary Information

**A. Dopamine depletion.** The effects of dopamine depletion in the BG network model have been replicated from the original study (27). A continuous parameter $\xi$ ranging from $0$ (physiological dopamine level) to $-0.8$ (low dopamine level) is included in the parameters, controlling some characteristic changes in network activity related to dopaminergic effects. Moreover, a scaling coefficient for the corresponding parameter fitted from experiments, which determines the relationship between dopamine receptor occupancy and the magnitude of the effect, is included (Table 8).

**B. Genetic algorithm.** The genetic algorithm was implemented with the PyGAD library (57), with the following settings:

- number of generations (i.e., the number of iterations that the genetic algorithm will run for): 5;
- number of parents (i.e., the number of solutions that will be selected for mating and producing offspring in each generation): 2;
- population size (i.e., the number of candidate solutions that will be evaluated in each generation): 2;
- number of genes (i.e., the number of parameters that make up each candidate solution): 4;
- parent selection method: steady-state selection;
- mutation type: random (i.e., each gene in a candidate solution has a probability of being randomly mutated);
- genes to mutate: 50 % ;
- crossover: single point;
- no stopping criteria.

| Population | Numerosity |
|---|---|
| Glomerulus (Glom) | 7073 |
| Granule cell (GrC) | 88164 |
| Golgi cell (GoC) | 219 |
| Molecular interneurons (MLI) | 1206 |
| Purkinje cell (PC) | 69 |
| Deep cerebellar nuclei (glutamatergic) (DCNp) | 12 |
| Deep cerebellar nuclei (gabaergic) (DNCi) | 12 |
| Inferior olivary cell (IO) | 12 |

Table 1: Cerebellar Numerosity

| Connection | Convergence (mean ± SD) | Divergence (mean ± SD) | $g$ [nS] | Delay [ms] | $\tau_\alpha$ [ms] | Type |
|---|---|---|---|---|---|---|
| Glom-GrC | 4 | 50 ± 22 | 0.15 | 4.0 | 5.8 | + |
| Glom-GoC | 65 ± 27 | 2 ± 1 | 1.5 | 4.0 | 0.23 | + |
| GoC-GrC | 2 ± 1 | 624 ± 267 | 0.6 | 2.0 | 13.6 | - |
| GoC-GoC | 34 ± 8 | 34 ± 9 | 0.3 | 1.0 | 10 | - |
| AA-GoC | 360 ± 81 | 1 | 1.2 | 2.0 | 0.5 | + |
| Pf-GoC | 1600 | 4 ± 2 | 0.05 | 5.0 | 0.5 | + |
| MLI-MLI | 4 ± 2 | 4 | 0.2 | 1.0 | 2 | - |
| PF-MLI | 1004 ± 221 (BC)<br>1021 ± 221 (SC) | 12 ± 4 (BC)<br>12 ± 5 (SC) | 0.015 | 5.0 | 0.64 | + |
| MLI-PC | 20 | 3 ± 1 | 0.3 | 4.0 (BC)<br>5.0 (SC) | 2.8 | - |
| AA-PC | 249 ± 13 | 1 | 0.7 | 2.0 | 1.1 | + |
| PF-PC | 28401 ± 776 | 23 ± 3 | 0.02** | 5.0 | 0.7 | + * |
| PC-DCNp | 26 ± 2 | 5 ± 1 | 0.4 | 4.0 | 1.14 | - |
| PC-DCNi | 26 ± 4 | 5 ± 1 | 0.12 | 4.0 | 1.14 | - |
| MF-DCNp | 147 | 1 | 0.05 | 4.0 | 1 | + |
| CF-PC | 1 | 6 ± 1 | 350 | 4.0 | 0.4 | + |
| CF-MLI | 3 ± 1 | 115 ± 23 | 1.0 | 70.0 ± 10.0 | 1.2 | + |
| IO-DCNp | 6 | 6 | 0.1 | 4.0 | 1 | + |
| IO_DCNi | 6 | 6 | 0.2 | 5.0 | 3.64 | + |
| DCNi-IO | 6 | 6 | 3.0 | 20. | 60.0 | - |

Table 2: **Cerebellar connections parameters**: $\tau_\alpha$ = time constants of the conductance, $g$ = conductance. Adapted from (26). Type is + for excitatory and - for inhibitory connections. * plastic synapse defined as in (40). ** adapted value to reproduce learning.

| | E-GLIF | | | | | | | | | | | | |
|---|---|---|---|---|---|---|---|---|---|---|---|---|---|
| Population | $C_m$ $[pF]$ | $\tau_m$ $[ms]$ | $E_L$ $[mV]$ | $t_{ref}$ $[ms]$ | $V_r$ $[mV]$ | $V_{th}$ $[mV]$ | $I_e$ $[pA]$ | $k_{adap}$ $[MH^{-1}]$ | $k_2$ $[ms^{-1}]$ | $A_2$ $[pA]$ | $k_1$ $[ms^{-1}]$ | $A_1$ $[pA]$ | $\lambda_0$, $\tau_V$ |
| GoC | 145 | 44 | -62 | 2 | -75 | -55 | 16.214 | 0.217 | 0.023 | 178.01 | 0.031 | 259.988 | 1, 0.4 |
| GrC | 7 | 24.15 | -62 | 1.5 | -70 | -41 | -0.888 | 0.022 | 0.041 | -0.94 | 0.311 | 0.01 | 1, 0.3 |
| MLI | 14.6 | 9.125 | -68 | - | -78 | -53 | 3.711 | 2.025 | 1.096 | 5.863 | 1.887 | 5.953 | 1.8, 1.1 |
| PC | 334 | 47 | -59 | 0.5 | -69.0 | -43 | 742.534 | 1.491 | 0.041 | 172.622 | 0.195 | 157.622 | 4, 3.5 |
| DCNp | 142 | 33 | -45 | 1.5 | -55 | -36 | 75.385 | 0.408 | 0.047 | 3.477 | 0.697 | 13.857 | 3.5, 3 |
| DCNi | 56 | 56 | -40 | 3.02 | -55 | -39 | 2.384 | 0.079 | 0.044 | 176.358 | 0.041 | 176.358 | 0.9, 1 |
| IO | 189 | 11 | -45 | 1 | -45 | -35 | -18.101 | 1.928 | 0.091 | 1358.197 | 0.191 | 1810.923 | 1.2, 0.8 |

Table 3: **Extended generalized leaky integrate and fire parameters**: $C_m$ = membrane capacitance; $\tau_m$ = membrane time constant; $E_L$ = resting potential; $t_{ref}$ = refractory period; $V_r$ = reset potential; $V_{th}$ = threshold potential; $I_e$ = endogenous current; $k_{adap}$, $k_2$ = adaptation constants; $k_1$ = decay rate of the intrinsic depolarizing current; $A_2$, $A_1$ = model currents update constants; $l_0$, $\tau_V$ = escape rate parameters. As in (34).

| Population | Numerosity |
|---|---|
| Fast spiking neurons (FSN) | 1599 |
| Medium spiny neurons with D1 receptors (MSN D1) | 37971 |
| Medium spiny neurons with D2 receptors(MSN D2) | 37971 |
| Globus pallidus externa type I (GPe TI) | 988 |
| Globus pallidus externa type A (GPe TA) | 329 |
| Subthalamic Nucleus (STN) | 388 |
| Substantia nigra pars compacta (SNr) | 754 |

Table 4: Basal Ganglia Numerosity

| | Delay [ms] | $E_{rev}$ $[mV]$ | $\tau$ $[ms]$ | $g$ $[nS]$ | Type | Synapse |
|---|---|---|---|---|---|---|
| FSN-FSN | 1.7 | -74 | 6 | 1 | - | Tsodyks |
| GPe-FSN | 7.0 | -74 | 17 (TI)<br>66 (TA) | 2 (TI)<br>0.51 (TA) | - | Static |
| GPe-GPe | 1 | -65 | 5 | 1.3 (TI)<br>0.33 (TA) | - | Static |
| MSN-GPe | 7 | -65 | 6 | 2 | - | Static |
| STN-GPe | 2 (TI) | 0 (TI) | 12 (TI) | 0.35 (TI)<br>0.11 (TA) | + | Static |
| MSN-MSN | 1.7 | -74 | 8 | 0.15 (D1-D1)<br>0.375 (D1-D2)<br>0.45 (D2-D1)<br>0.35 (D2-D2) | - | Static |
| FSN-MSN | 1.7 | -74 | 11 | 6 | - | Tsodyks |
| GPe-MSN | 7.0 | -74 | 87 (TA-D1)<br>76 (TA-D2) | 0.04 (D1)<br>0.08 (D2) | - | Static |
| MSN-SNr | 7.0 | -80 | 5.2 | 2 | - | Tsodyks |
| GPe-SNr | 3 | -72 | 2.1 | 76 | - | Tsodyks |
| STN-SNr | 4.5 | 0 | 12 | 0.91 | + | Tsodyks |
| GPe-STN | 1 | -84 | 8 | 0.08 | - | Tsodyks |

Table 5: **BG connections parameters**: $E_{rev}$ = reversal potential; $\tau$ = time constants of the conductance, $g$ = conductance. As in (27).

| | U | $\tau_{rec}$ $[ms]$ | $\tau_{fac}$ $[ms]$ |
|---|---|---|---|
| FSN-MSN | 0.29 | 902 | 53 |
| FSN-FSN | 0.29 | 902 | 53 |
| GPe-SNr | 0.196 | 969 | 0 |
| MSN-SNr | 0.0192 (D1)<br>0.24 (D2) | 623 (D1)<br>11 (D2) | 559 (D1)<br>73 (D2) |
| STN-SNr | 0.35 | 800 | 0 |
| GPe-STN | 0.29 (TI)<br>0.29 (TA) | 902 (TI)<br>902 (TA) | 53 (TI)<br>53 (TA) |

Table 6: **BG Tsodyks synapse parameters**: $U$ = utilization factor ; $\tau_{rec}$ = recovery time constant, $\tau_{fac}$ = facilitation time constant. As in (27).

Izhikevich

| Population | $a$ $[s]$ | $\alpha$ | $b$ | $c$ $[mV]$ | $C_m$ $[pF]$ | $d$ $[pA]$ | $k$ | $\nu_{peak}$ $[mV]$ | $E_L$ $[mV]$ | $V_{th}$ $[mV]$ | $\nu_b$ $[mV]$ |
|---|---|---|---|---|---|---|---|---|---|---|---|
| FSN | 0.2 | - | 0.025 | -60 | 80 | 0 | 1 | 25 | -64.4 | -50 | -55 |
| MSN | 0.01 | 0.032 | -20 | -60 | 15.2 | 66.9 (D1)<br>91 (D2) | 1 | 40 | -78.2 (D1)<br>-80 (D2) | -29.7 | - |

Adaptive exponential integrate and fire neuron

| Population | $a$ $[nS]$ | $b$ $[pA]$ | $C_m$ $[pF]$ | $\Delta_T$ $[ms]$ | $E_L$ $[mV]$ | $g_L$ $[nS]$ | $I_e$ $[pA]$ | $\tau_w$ $[ms]$ | $t^f$ $[mV]$ | $V_r$ $[mV]$ | $V_T$ $[mV]$ |
|---|---|---|---|---|---|---|---|---|---|---|---|
| GPe | 2.5 | 70 (TI)<br>105 (TA) | 40 (TI)<br>60 (TA) | 1.7 (TI)<br>2.55 (TA) | -55.1 | 1 | 12 (TI)<br>1 (TA) | 20 | 15 | -60 | -54.7 |
| SNr | 3 | 200 | 80 | 1.8 | -55.8 | 3 | 15 | 20 | 20 | -65 | -55.2 |
| STN | 0.3 | 0.05 | 60 | 16.2 | -80.2 | 10 | 5 | 333 | 15 | -70 | -64.0 |

Table 7: **Izhikevich neuron parameters** $a$ = Recovery current time constant; $b$ = Spike-triggered adaptation; $\beta_{E_L}$ = Magnitude of D1 effect on resting potential; $C_m$ = membrane capacitance; $\Delta_T$ = Slope factor of spike upstroke; $E_L$ = Resting potential; $g_L$ = Leak conductance; $I_e$ = current injected to have desired fr ($8Hz$ for TA, $18Hz$ for TI); $t^f$ = Spike cutoff; $V_r$ = reset potential. **Adaptive exponential integrate and fire neuron parameters** $a$ = Subthreshold adaptation; $b$ = Voltage dependency of recovery current; $c$ = Spike reset; $C_m$ = membrane capacitance; $d$ = Summed recovery current contribution following an action potential; $k$ = Steady-state voltage dependence; $\nu_b$ = Voltage dependence recovery current; $\nu_{peak}$ = Spike cutoff; $v_r$ = Resting potential; $V_{th}$ = Threshold potential; $\tau_w$ = Adaptation time constant. As in (27)

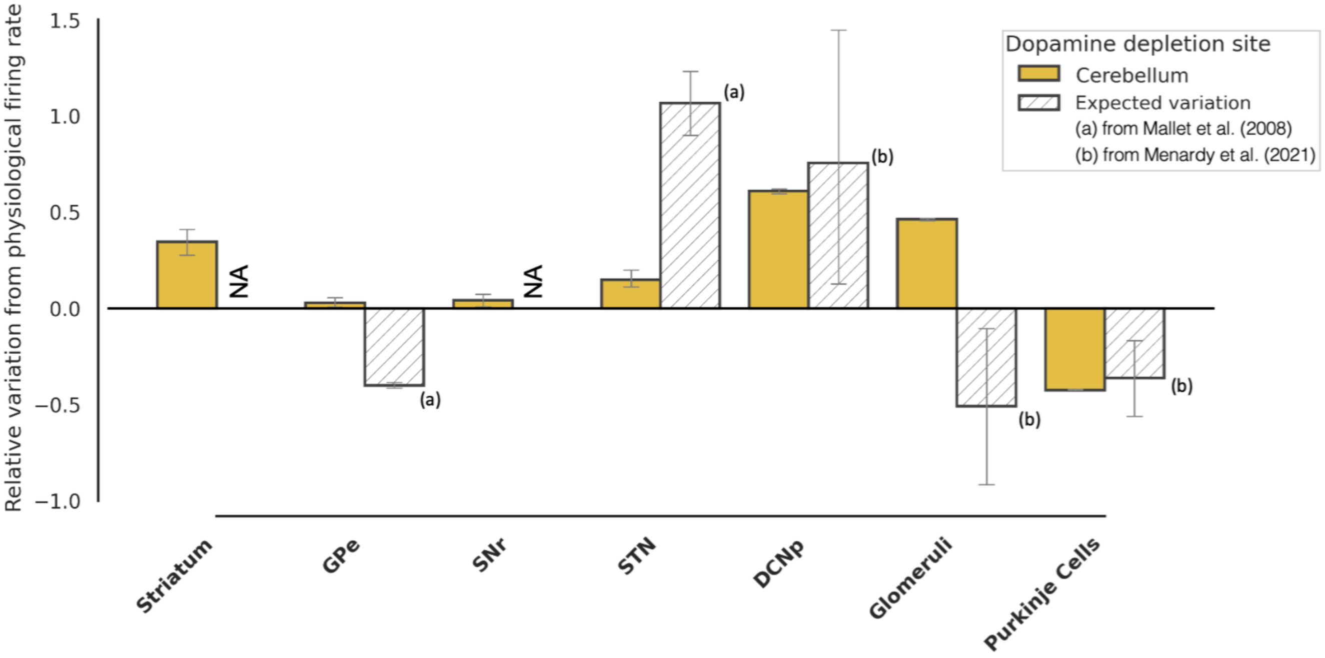


**Fig. 6. Firing rate variations with dopamine depletion in only Cerebellum**: histograms of normalized firing rate variations in the main spiking populations. The reported values are computed considering the most severe pathological scenario (depletion level = 0.8), as values reported from literature refer to advanced pathology (6, 49) and have been obtained by normalizing the difference between the firing rates in pathological and physiological conditions with respect to the physiological firing rate ($\Delta_{fr} = \frac{fr_{pathological} - fr_{physiological}}{fr_{physiological}}$); therefore, a value of 0 corresponds to no difference between the two, while a value of 1 corresponds to a pathological firing rate that is two times the physiological one.

| | | **Parameter ($p$)** | **$\beta$ factor** | **Rule** |
|---|---|---|---|---|
| **Neuron model** | FSN | $\nu_r$ | -0.078 | $p(1+\beta\xi)$ |
| | GPe | $E_L$ | -0.181 | $p(1+\beta\xi)$ |
| | MSN-D1 | $\nu_r$ | 0.0296 | $p(1+\beta\xi)$ |
| | MSN-D1 | $d$ | -0.450 | $p(1+\beta\xi)$ |
| | SNr | $E_L$ | -0.0896 | $p(1+\beta\xi)$ |
| **Connection model** | FSN-FSN | $I_{GABA}$ | -1.27 | $p(1+\beta\xi)$ |
| | GPe-FSN | $I_{GABA}$ | -0.53 | $p(1+\beta\xi)$ |
| | GPe-GPe | $I_{GABA}$ | -0.83 | $p(1+\beta\xi)$ |
| | MSN D2-GPe TI | $I_{GABA}$ | -0.83 | $p(1+\beta\xi)$ |
| | STN-GPe | $I_{AMPA}$ | -0.45 | $p(1+\beta\xi)$ |
| | CTX-MSN D1 | $I_{NMDA}$ | 1.04 | $p(1+\beta\xi)$ |
| | CTX-MSN D2 | $I_{AMPA}$ | -0.26 | $p(1+\beta\xi)$ |
| | MSN-MSN | $I_{GABA}$ | 0.88 | $p(1+\beta\xi)$ |
| | GPe TI-MSN D1 | $I_{GABA}$ | -1.22 | $p(1+\beta\xi)$ |
| | GPe TA-MSN D2 | $I_{GABA}$ | -1.15 | $p(1+\beta\xi)$ |
| | MSN-SNr | $I_{GABA}$ | 0.56 | $p(1-\beta\xi)$ |
| | CTX-STN | $I_{GABA}$ | -0.45 | $p(1-\beta\xi)$ |
| | GPe-STN | $I_{GABA}$ | -0.24 | $p(1-\beta\xi)$ |

Table 8: **Multiplicative factor controlling effects of dopamine depletion**: $v_r$, $E_L$ = resting potential; $d$ = Summed recovery current contribution following an action potential; $I_{GABA}$, $I_{NMDA}$, $I_{AMPA}$ = GABA, NMDA, AMPA currents (defined as $I = g(E_{rev} - V)$, with $g$, $E_{rev}$ and $V$ being the conductance, the reversal potential and the membrane potential correspondingly)

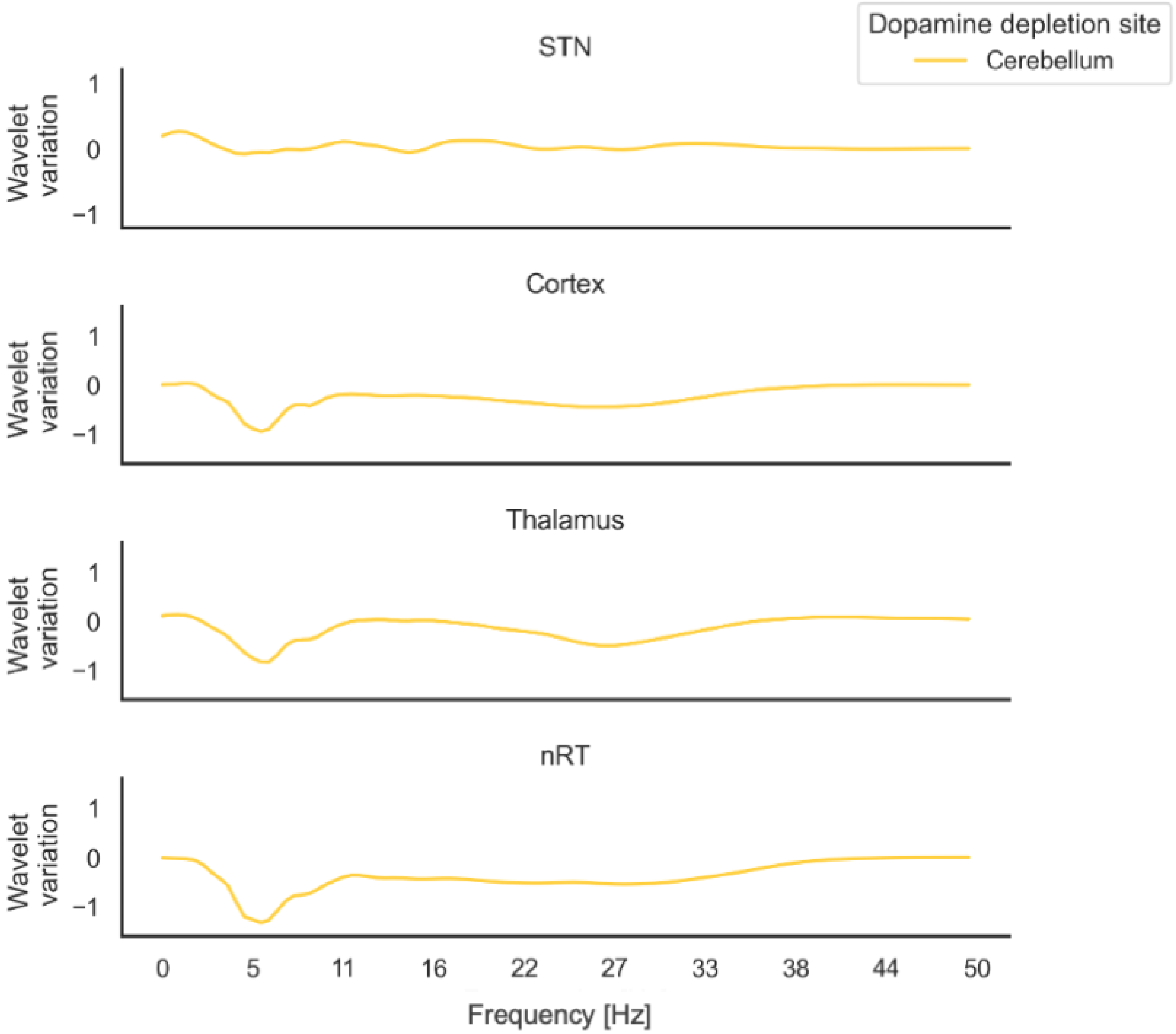


**Fig. 7. Spectral variation with dopamine depletion in Cerebellum**: spectrum values of the three mass models (Cortex, Thalamus, nRT) and subthalamic nucleus (STN). Each curve represents the variation of the power spectrum evaluated at the most severe case (depletion level -0.8) with respect to the physiological condition ($\Delta_{W_T} = \frac{W_T\ pathological - W_T\ physiological}{W_T\ physiological}$).

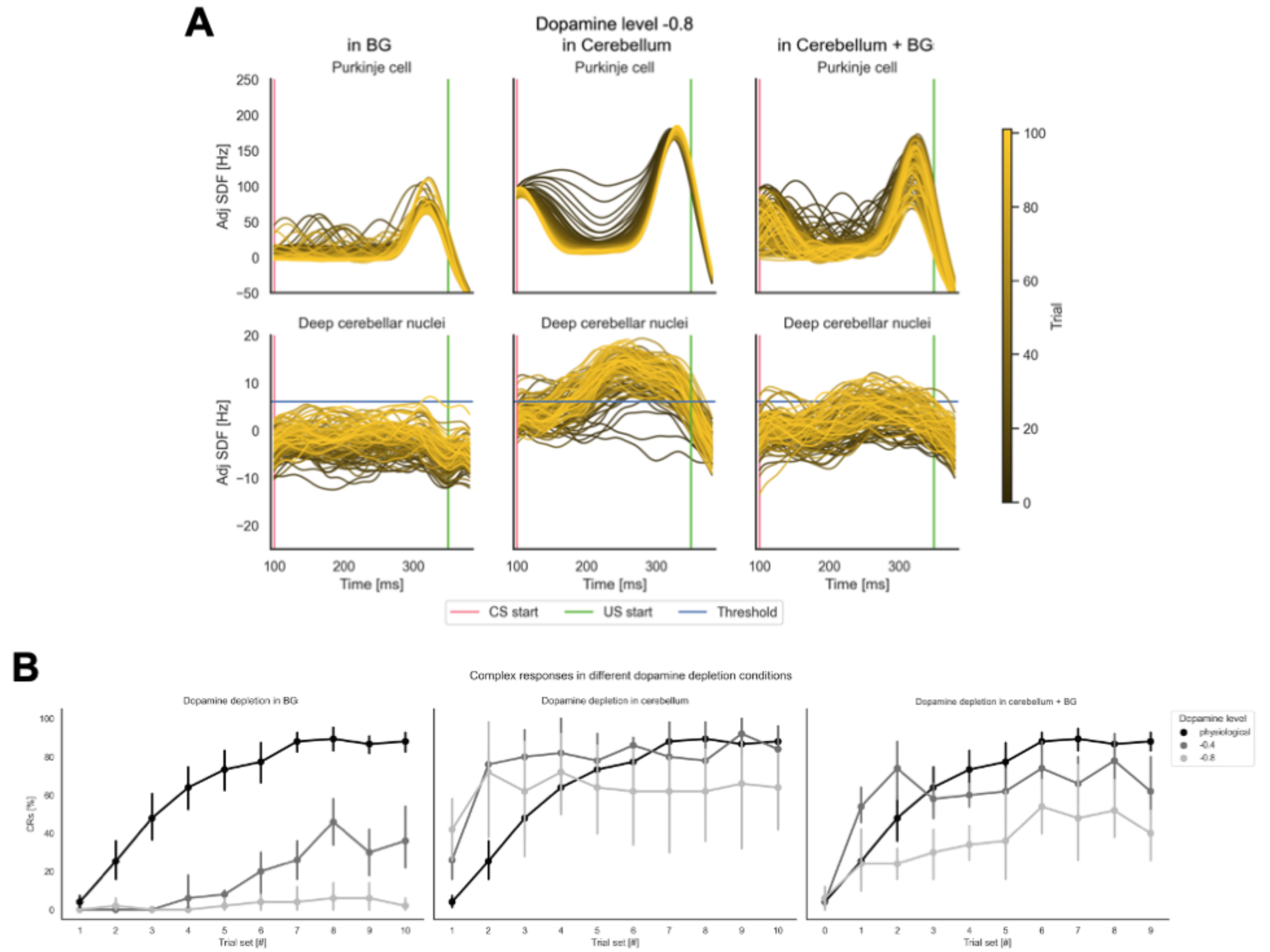


**Fig. 8. Learning protocol**: eye-blink classical conditioning results both in terms of activities of the main populations of interest (PCs and DCNp) and in terms of %CR. Panel **A** shows the results of the EBCC protocol when dopamine depletion is modeled in different sites (only BG on the left, only Cerebellum in the center, Cerebellum + BG on the right) at the most severe case (depletion level -0.8). The entire CS window is shown, with all trials overlaid (each curve represents the SDF within a single trial, and trials' progression is shown with a shaded color). The signals are adjusted for the baseline evaluated in the 200 ms following the stimuli. Unconditioned stimulus (US) start and threshold for identification of CR are reported. In panel **B**, the percentage of CR identified in each 10-trial block, at different sites (as in panel **A**) and magnitudes of dopamine depletion (0 = physiological state, -0.4, -0.8) are overlaid.

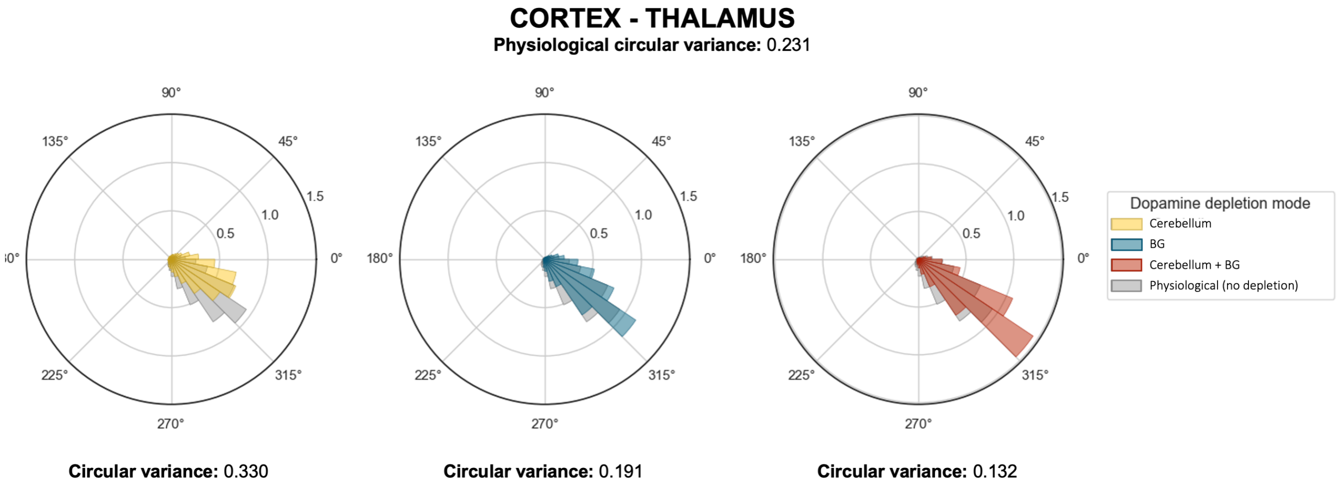


**Fig. 9. Circular variance of mass models' activities**: polar histograms of the phase differences between mass models' activities and the corresponding circular variance values in three different sites of dopamine depletion (Cerebellum, BG, and Cerebellum + BG, with depletion level $\xi = -0.8$ and in physiological conditions).
